# Rationalizing sensorimotor deficits in autism spectrum disorder

**DOI:** 10.64898/2026.10.12.758517

**Authors:** Yizhou Chen, Jean-Paul Noel, Paul Schrater, Dora Angelaki, Xaq Pitkow

## Abstract

Autism spectrum disorder (ASD) is a neurodevelopmental condition characterized by socio-communicative, sensory, and motor anomalies. Recently, a host of probabilistic models have been proposed to account for the varied phenotypes observed in ASD. These computational psychiatry efforts, however, have (1) largely focused on simple tasks that artificially segregate action from perception, and (2) unequivocally assumed that both neurotypical and divergent populations attempt to optimize the same cost function. Here, we ask individuals with ASD and neurotypical (NT) controls to perform a task in virtual reality (VR): they should navigate to, and stop at, the position of briefly presented targets in a closed action-perception loop. First, we demonstrate that while summary task performance is insufficient to distinguish between ASD and NT individuals, data-rich virtual trajectories do differentiate these groups. This suggests that ASD and control participants use distinct navigational strategies and/or cost functions. Next, we leverage the theoretical framework of Inverse Rational Control (IRC) to interpret the sensorimotor anomalies present in ASD. IRC does not assume that agents are optimal, but rational: that is, agents attempt to maximize reward according to an internal model of the task, which may not match the true task and environment. Applying IRC to this VR navigation task revealed both sensory and motor differences between ASD and neurotypical participants. The sensory differences are consistent with Bayesian inference under altered weighting of sensory evidence. The motor differences manifest as more frequent and abrupt actions in ASD, which we explain by a reduced subjective action cost in ASD. Together, these results demonstrate the utility of model-based characterization of psychopathology and suggest that the computational study of ASD should consider not only perceptual anomalies, but also subjective costs.

## Introduction

Autism spectrum disorder (ASD) is a heterogenous neurodevelopmental condition afflicting a wide range of socio-communicative (***American Psychiatric Association, 2013***), perceptual (***Robertson and Baron-Cohen, 2017***), and motor (***Fournier et al., 2010***) functions. To account for these varied phenotypes with a unified explanation, researchers have cast the disorder in the language of probabilistic inference, characterized by an imbalance in the relative weighting of predictions from Bayesian priors (***Van de Cruys et al., 2014***; ***Lawson et al., 2017***; ***Lieder et al., 2019***; ***Vishne et al., 2021***; ***Noel et al., 2024a***) against sensory observations (i.e., Bayesian likelihoods; ***Pellicano and Burr (2012***)). However, the experimental studies of ASD have largely used relatively simple tasks that under-constrain model selection. Namely, typical psychophysical tasks aimed at understanding perceptual processing in ASD often require only intermittent binary responses, and artificially segregate action from perception (but see ***Arthur et al. (2021***) for a recent exception). Instead, in the real world, action and perception are not independent.

The probabilistic modeling accounts of ASD have followed suit, largely ignoring the motor system by studying Bayesian inference but not optimal control, and assuming that ASD and neurotypical individuals attempt to optimize the same objective. Humans, however, are often not optimal under many circumstances (***Beck et al., 2012***), and is it not clear that neurotypical individuals and neurodivergent ones (e.g., those with ASD) even share a cost function.

Here, we address the above experimental and computational limitations by measuring behavior during a task that requires participants to continually act upon a latent variable – their distance to a hidden target – in a closed action-perception loop. Participants see a virtual environment composed solely of flashing ground-plane textural elements, and must navigate via path integration to the location of a briefly presented target (see ***Lakshminarasimhan et al. (2018***); ***Noel et al. (2020***, 2022, 2024b); ***Alefantis et al. (2022***)). As in real-world behavior, this task requires continual updating of beliefs about task-relevant variables: in this case, about the relative displacement between self and target.

To analyze these behavioral data, we reverse engineer each participant’s internal model of the task dynamics under the assumption that they are *rational*, rather than *optimal*. That is, we hypothesize that agents attempt to maximize reward/minimize cost within a potentially incorrect internal model of the world (***Wu et al., 2020***; ***Kwon et al., 2020***). We fit full trajectories using the theoretical framework of Inverse Rational Control (IRC; ***Wu et al. (2020***); ***Kwon et al. (2020***)) which estimates the participants’ internal model, reward/cost function, and dynamic beliefs. This approach substantially expands on previous probabilistic frameworks applied to ASD, as well as previous attempts to account for navigation toward hidden targets (***Lakshminarasimhan et al., 2018***; ***Noel et al., 2020***) by modeling not only the perceptual inference process, but also dynamic control and subjective value.

We demonstrate that while neurotypical participants and individuals with ASD had the same mean task performance, they had different navigational trajectories and task strategies. We use IRC to interpret these distinct strategies. The internal models of individuals with ASD reveal different assumptions about how the world works: they weigh their sensory observations less, and correct their errors using actions that are stronger but subjectively cheaper. These two effects explain the more frequent and abrupt actions in ASD, highlighting the utility of modeling both action and perception.

## Results

### Distinct navigational strategies with similar outcomes

Individuals with ASD (*n* = 14) and their neurotypical counterparts (NT, *n* = 25) were tasked with “catching fireflies” in virtual reality. Participants used a joystick to control their virtual velocity in two dimensions (forward and angular) and move within the environment before stopping at the target location. The environment had no fixed landmarks and instead was composed solely of transient ground plane textural elements (***Figure 1*A**). As participants navigated, the ground plane elements correspondingly moved on their retina, generating an optic flow signal informative about their self-motion velocity. To estimate position from this evidence, participants had to continuously integrate the visual velocity signal (***Figure 1*A, B**) into an estimate of position. Importantly, although not incorporated into previous modeling efforts (e.g., ***Noel et al. (2020***)), participants implicitly had access to a second source of velocity information: the velocity predicted from their internal model of the joystick gain together with an efference copy of their joystick movements. We consider the velocity information provided by optic flow to be their “observations,” leading to a likelihood in the probabilistic framework; the knowledge available from their internal model and efference motor copy provides a “prediction” of the velocity, which we consider a Bayesian prior.

**Figure 1.**
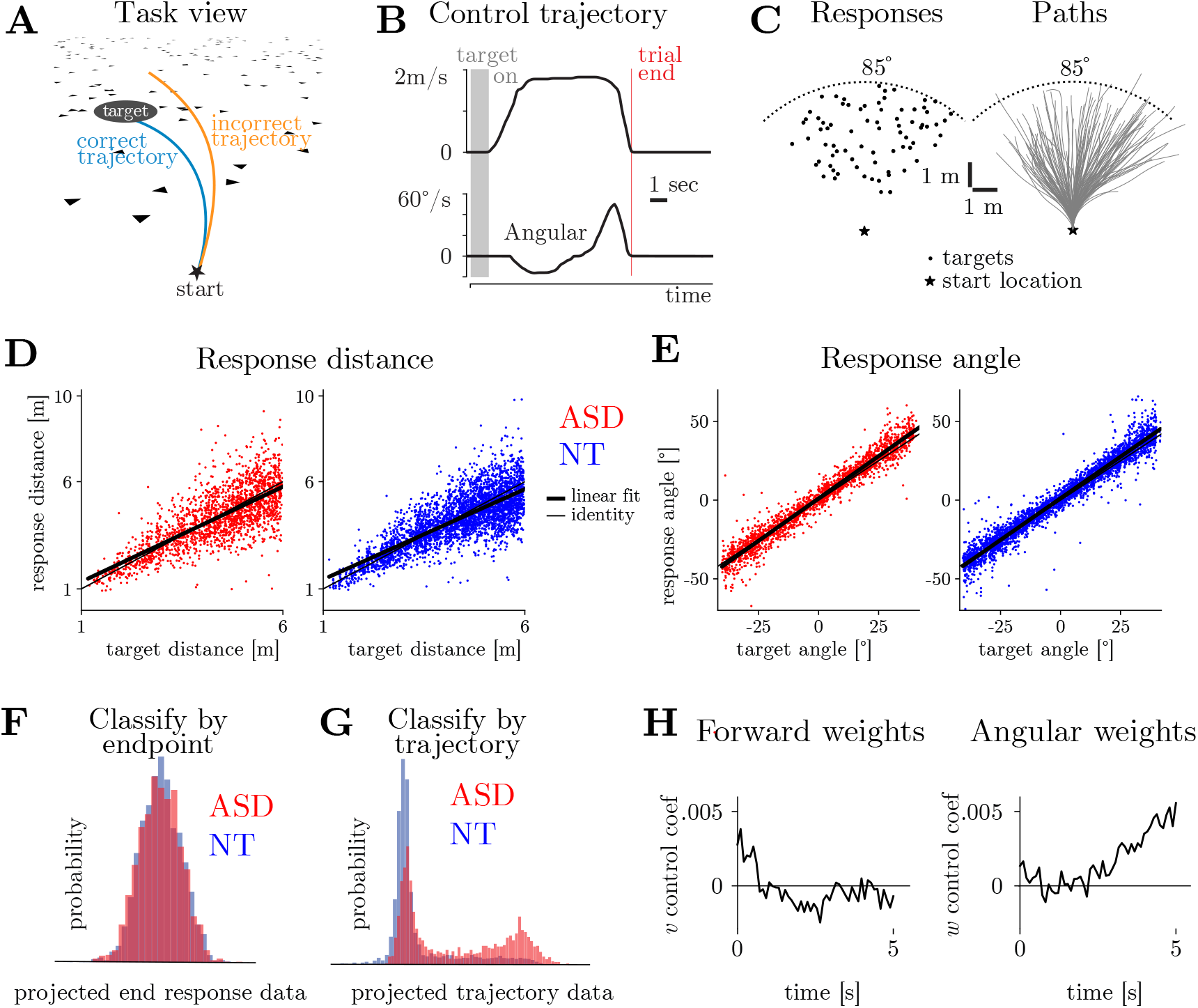
The firefly task and model-free analysis of behavior. **A**. Task schematic. The target (black circle) is visible at the beginning of the trial (1s), and participants use a joystick and the optic flow provided by ground plane textural elements (black triangles) to navigate and stop at the target location. **B**. Velocity profiles in the forward (top) and angular (bottom) dimension for an example trial from a control subject. The black shaded area corresponds to the time the target is visible. Red vertical line is the end of the trial. **C**. A different target location is presented on each trial, within a distance range from 1 to 6 m, and −42.5 to 42.5 degrees. *Left*: Target locations for an example subject. *Right*: Trajectories toward the presented targets. **D-E**. Endpoint locations in the forward (D) and angular (E) dimension as a function of target location for all individuals with ASD (red) and neurotypical controls (blue). Thin lines have a slope of 1, showing no bias (slope = 1). Thick lines show best-fitting lines. Best fitting slopes for response distance: ASD = 0.89, NT = 0.84; for response angle: ASD = 1.08, NT = 1.06. **F**. The distribution of ASD (red) and NT (blue) endpoint responses projected onto the most separable axis according to a linear Support Vector Machine (SVM). The SVM was not able to distinguish between the two groups. **G**. The distribution of ASD and NT trajectory data projected onto the most separable axis as estimated by SVM. SVM was able to distinguish the two groups. **H**. The SVM weights of the forward and angular velocity separate the two groups as a function of time.

In each trial, a target “firefly” was presented at a random location for 1s within a radial distance of 1 to 6 meters, and angular eccentricity between ± 42.5 degrees from straight ahead (***Figure 1*C**). Upon stopping, participants received feedback about their performance by a tone and an arrow pointing to the center of the target. The arrow was colored green for correct trials, designated when participants stopped within 65 cm of the target, and red for incorrect ones. Before the experiment, participants were familiarized with the task and virtual environment over *∼*150 trials.

As a first approach to quantifying task performance, we decomposed trajectory endpoints and target positions into their forward and angular components (see ***Lakshminarasimhan et al. (2018***); ***Noel et al. (2020***) for a similar approach). ***Figure 1*D** and **E** shows that the relation between trajectory endpoints and targets was approximately linear (slope = 1 indicates no bias; mean ± SEM; ASD *R*^2^ = 0.93 ± 0.08; NT *R*^2^ = 0.85 ± 0.05; *t*-test comparing group: forward *p* = 0.53; angular *p* = 0.59). Overall, both groups overshot targets in the angular dimension (slope = 1 indicates no bias; slope for ASD = 1.08 ± 0.02; slope for NT = 1.06 ± 0.02; All *t*-test *p* < 0.05), and undershot them in the forward dimension (slope for ASD = 0.89 ± 0.04; slope for NT = 0.84 ± 0.02; all *t*-test *p* < 0.05). There was no significant difference in the trajectory endpoints for individuals with ASD versus neurotypical controls (both *t*-tests had *p* > 0.50). Both ASD and NT participants showed no difference in performance early versus late in the session, indicating a stable behavior (both *t*-tests had *p* > 0.10; ***Appendix 1 Figure 2***).

We could not distinguish between ASD or NT individuals by their trajectory endpoints (classification by linear Support Vector Machine (SVM) (***Cortes and Vapnik, 1995***): Kolmogorov-Smirnov test, *p* = 0.80); ***Figure 1*F**). However, an SVM did reliably distinguish these groups based on their full navigation trajectories (linear SVM on time series, zero-padded to a fixed length; Kolmogorov-Smirnov test had *p* < 0.001). As shown in ***Figure 1*G**, the projected ASD trajectory data showed a stereotyped second peak that was much less prominent in the NT data. This effect was preserved at the single participant level (***Appendix 1 Figure 3***). The time course of SVM coefficients indicated which task variables best distinguished the experimental groups, and when: ASD participants had higher early forward velocities and late angular velocities compared to the NT participants (***Figure 1*H**). These differences result in the stereotyped second peak seen in ***Figure 1*G**.

Although this data-driven approach can show that ASD and NT use different navigational strategies, it cannot interpret or describe *why* these differences exist. To address this question, in the next section, we use a model-based approach to reverse engineer the navigational strategies used by ASD and NT participants.

**Table 1.** Task-relevant parameters included as components of *θ*. Each entry has both forward (*v*) and angular (*w*) components. Action cost terms are expressed in units of state rewards.

| Subjective parameter $\theta$ | Explanations |
| --- | --- |
| Process gain | Scale factor mapping joystick position to virtual velocity. |
| Process noise | Standard deviation of system's process noise, *i.e.* variability added to world state. |
| Observation noise | Standard deviation of participant's observation noise. |
| Initial uncertainty | Standard deviation of participant's initial uncertainty about target. |
| Action cost (derivative) | Scale factor for computing subjective cost from Total Squared Variation (TSV) of the joystick deflection. |
| Action cost (magnitude) | Scale factor for computing subjective action cost from squared joystick deflection from origin. |
| Objective parameter $\phi$ | Explanations |
| Process gain | Scale factor mapping joystick position to virtual velocity. |
| Process noise | Standard deviation of system's actual process noise. |

### Inverse Rational Control as a framework accounting for behavior in closed action-perception loops

We argue that humans are rational (***Wu et al., 2020***). That is, instead of adopting policies that would objectively maximize reward (i.e., optimal behavior), they may adopt policies maximizing a subjective reward function based on (perhaps erroneous) assumptions about the world dynamics. To allow for this possibility, we estimate an agent model of each participant, using Inverse Rational Control (IRC) (***Wu et al., 2020***; ***Kwon et al., 2020***). IRC is grounded in a mathematical model of the task behavior called a Partially Observable Markov Decision Process (POMDP; ***Figure 2*A**, ***Spaan (2012***)). This process determines how optimal participants should navigate to the location of the hidden fireflies. We characterize the firefly task by task variables that depend on time *t*, and by parameters that are independent of time. The variables include the world state *s*_*t*_ comprising the relative position, heading, and velocity of the observer compared to the firefly target, the joystick controls or actions *a*_*t*_, and sensory observations given by optic flow *o*_*t*_. Some of the parameters describe the dynamics of the task: how actions affect the forward and angular velocities in the world (“process gains”), how much variance is added to these velocities (“process noise” variances), how strong the mean optic flow observations are for given self-motion velocities (observation gains), and how informative the sensory data is (observation noise variances). Additional parameters describe the instantaneous rewards (rewarded region and reward amounts) and the cost of moving, which we denote together as *R*_*t*_(*s*_*t*_, *a*_*t*_ *θ*). Another parameter describes a temporal discount factor *γ* which limits how long the agent plans into the future. We denote by *θ* the vector of all parameters together (***Appendix 1 Table 1***). Although our rational model assumes that the participant knows the true *structure* of the task (the relevant variables, which variables interact, etc.), it allows that the participant may assume different parameters *θ* than the true experimental parameters *ϕ*, such as believing that the joystick gain is lower than it really is experimentally (***Wu et al., 2020***; ***Kwon et al., 2020***; ***Stocker and Simoncelli, 2006***; ***Lakshminarasimhan et al., 2018***). Additionally, some of the parameters are purely subjective and have no objective truth, like the worth of a firefly compared to the cost of catching it.

**Figure 2.**
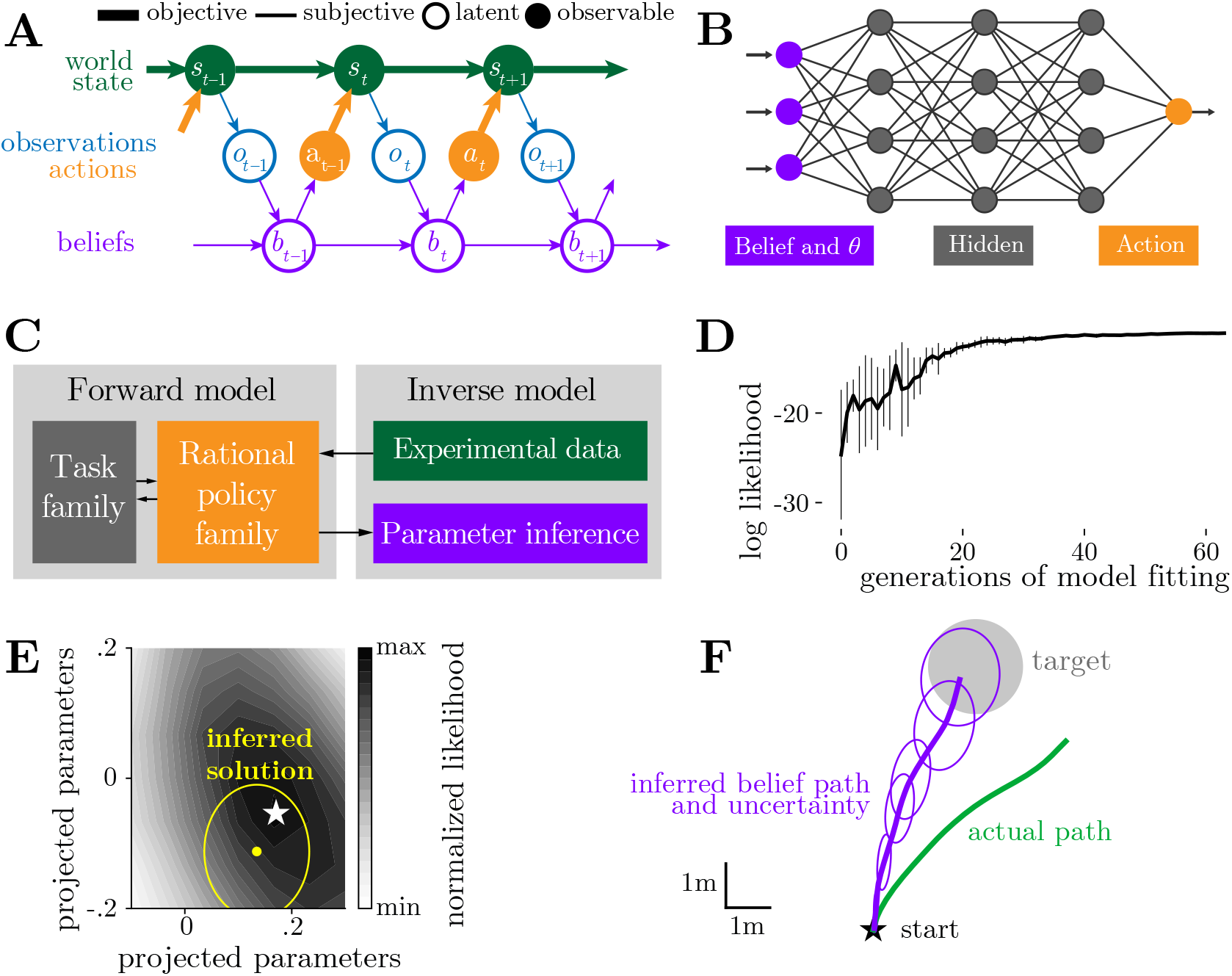
Inverse rational control (IRC) framework. **A**. We model the task as a Partially Observable Markov Decision Process (POMDP) wherein participants choose actions (orange) to maximize their long-term subjective value, given their internal model of the task. World state (*s*, green) describes the current state of the task, including the virtual participant’s position and velocity. The participant takes an observation (*o*, blue) of the world state, and integrates it with the previous belief (*b*_*t*−1_, purple) about the world state to form the new belief (*b*_*t*_, purple) about the current world state. Finally, the participant’s action (*a*, orange) is to control the joystick, which changes their world state by moving towards the target state (goal of the task). **B**. An agent’s policy, or probability of each action given the current belief and task parameters, is parameterized by a feedforward neural network. **C**. The IRC framework has two components, a forward and an inverse model. In the forward model, we use reinforcement learning to find approximately optimal policies for solving a family of tasks. In the inverse model, we identify which of these forward models best accounts for behavioral data. **D**. The observable data log-likelihood is maximized during the inverse fitting process in evaluation trials. Error bars indicate one standard deviation over trials after each training epoch. **E**. The inverse algorithm finds an appropriate solution near the ground truth on simulation data. Colors indicate observed data likelihoods over a 2D slice of parameter space. White star: ground truth parameters *θ*. Yellow ellipse centered at yellow dot: inferred best-fitted parameters 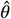 with uncertainty (1 standard deviation). **F**. Overhead view of a subject’s trajectory (green) and its corresponding estimated belief trajectory (purple trajectory; ellipses represent the uncertainty of belief) given the inferred internal model of the subject. When the participant stops moving, it believes it has stopped on the target (grey disk) with some uncertainty, although the actual trajectory had missed the target.

Given these parameters, a POMDP agent uses Bayesian inference to synthesize evidence over time, weighing its partially reliable observations (the “likelihood”) against its actions (which, through an internal model, produce a dynamic “prior” distribution over the world state. (See Methods Inverse Rational Control framework.) This yields a posterior probability distribution over the hidden world state, *p*(*s*_*t*_ *o*_1:*t*_, *a*_1:*t*−1_, *θ*), which we describe as a belief *b*_*t*_. Based on this belief, the agent then chooses actions at each time according to a policy *π*(*a*|*b*_*t*_, *θ*) that maximizes its value, defined as the total expected (temporally-discounted) subjective long-term reward,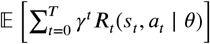.

We can fit this POMDP agent model to behavioral data. Our approach first finds optimal policies for a whole family of possible world dynamics that our participants might hypothesize and subjective rewards they might experience. We call this phase of the modeling approach “solving the forward model.” Concretely, we estimate the optimal policy using reinforcement learning (RL, ***Figure 2*B**), specifically the RL algorithm TD3 (***Fujimoto et al., 2018***).

Then, we find which of these optimal policies best accounts for the observed human behavior. We call this second phase “solving the inverse model.” Specifically, we maximize the experimentally observable data likelihood (Methods Inverse model: fitting the latent parameters) to find the parameters 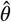 and corresponding policy *π* that best explain a participant’s behavior when catching fireflies. The whole process is depicted in ***Figure 2*C**, with more details in Methods Inverse Rational Control framework and ***Wu et al. (2020***); ***Kwon et al. (2020***).

We validate this IRC approach in simulated data (also see ***Wu et al. (2020***); ***Kwon et al. (2020***)). An artificial agent is trained to perform the firefly task governed by one set of subjective parameters *θ*, which we take to be the ground truth for that agent. Then, we test this agent on a task with a different set of experimental task parameters *ϕ*. We use an iterative search algorithm, CMAES (***Hansen (2023***), Methods Inverse model: fitting the latent parameters), to fit this behavior by solving the inverse problem, thereby inferring the agent’s maximum likelihood internal model 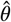 about the task. We observe that IRC converges onto a set of parameters that best explain the agents’ behavior (***Figure 2*D**). To visualize how IRC approaches the true parameters, we plotted the log-likelihood surface on a 2D slice defined by the two parameter dimensions that exhibited the greatest change during optimization. We also plotted the true parameters, final IRC estimate, and confidence region. ***Figure 2*E** demonstrates that the inverse algorithm finds a solution near the ground truth parameters (white star). ***Figure 2*F** plots the true target position (black circle) and the agent’s actual trajectory (green), as well as the agent’s belief, including both its mean estimate (thick line) and uncertainty (purple covariance ellipses). Even though the simulated agent did not correctly stop within the target, it *believes* it did: the belief trajectory ends inside the target, which rationalizes why the agent stopped.

### Applying IRC to data to discover reliable explanations of behavior

Having demonstrated the utility of the approach in synthetic data, we applied IRC to the trajectories taken by ASD and NT participants to identify which properties of their internal models best account for their distinct task strategies. In other words, we are making mathematical models of their mental models. The mathematical models accounted for ASD and NT behavior with comparable likelihood (mean difference in log-likelihood between training and testing data was −1.19; *t*-test training vs testing log-likelihood, *p* = 0.26; see Methods Validation of the IRC on simulation data and ***Appendix 1 Figure 5***). The fitted models generated trajectories (***Figure 3*A**) and velocity profiles (***Figure 3*B**) that recapitulated human behavior for both training and testing data. The inferred parameter set for each participant is shown in ***Figure 3*C**.

**Figure 3.**
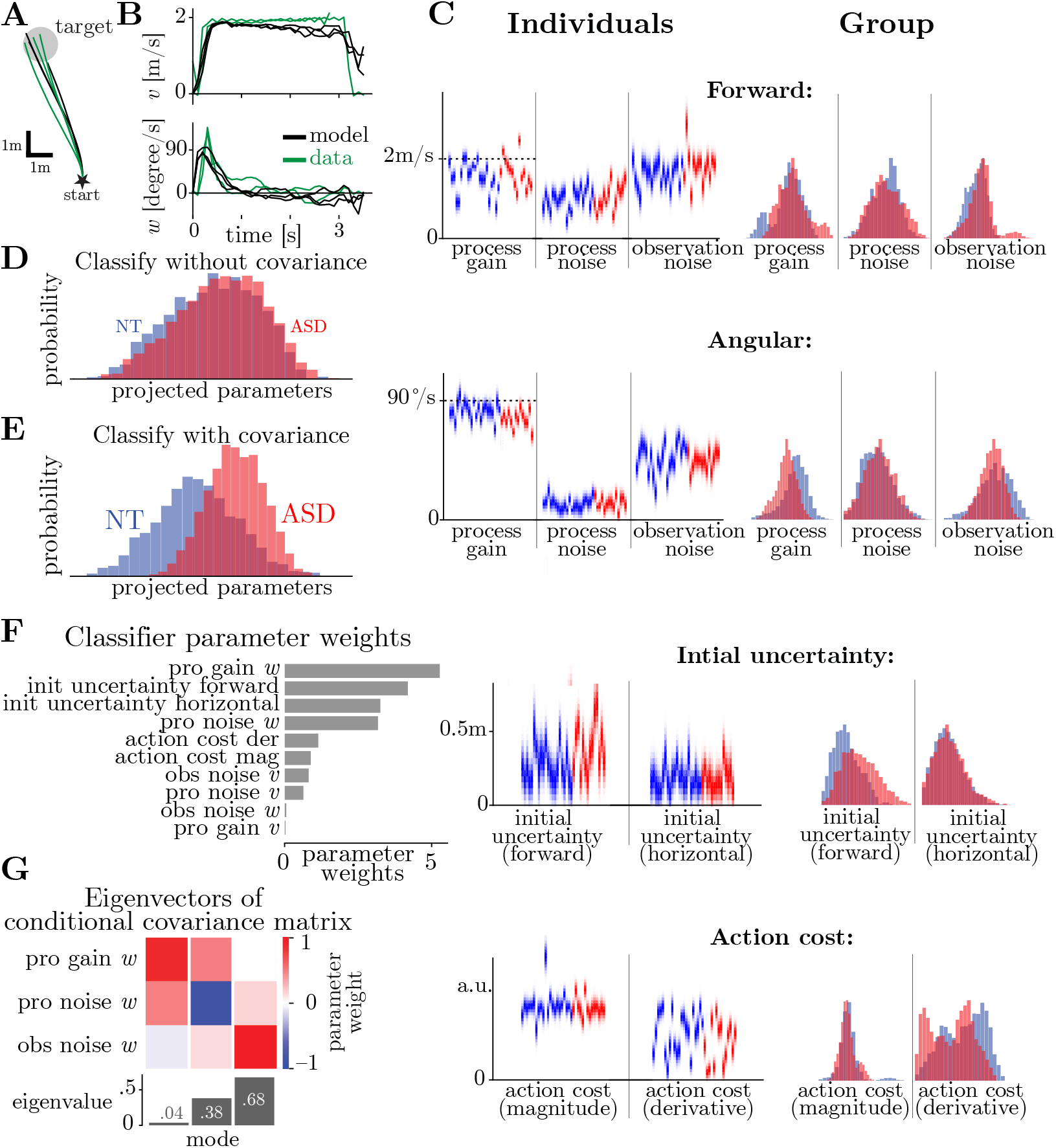
IRC infers participants’ internal models, and interprets behavioral differences between ASD and neurotypical participants. Trajectories (**A**) and velocity profiles (**B**) produced by IRC match those from human participants. **C**. Parameter estimates from IRC 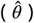 for ASD (red) and neurotypical (NT) participants (blue). Left: each pixel column indicates the estimated parameter vector of one subject, shaded by its uncertainty according to CMA-ES. Ground truth process gains are shown as dashed lines. Right: ASD and NT population parameter distributions. Refer to Methods ***Table 1*** for descriptions of parameters and Appendix 1 ***Table 3*** for specific values of inferred parameters. **D**. The distribution of the estimated assumptions and subjective preferences 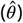 across participants, projected onto the axis that was best separated by a Support Vector Machine (SVM). This SVM cannot distinguish the ASD and NT groups without accounting for the *joint* likelihood between parameters. **E**. Distribution of the subject’s parameters estimated by IRC 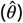 projected onto the most separable axis solved by an SVM. The two groups are much more separable when given the covariance information between each pair of parameters. **F**. The absolute SVM coefficients for the inferred parameter estimates in Panel D. The parameters that best distinguish the two groups are the estimated assumed gain, estimated assumed process noise, and derivative action cost. **G**. Eigenvectors and eigenvalues of the covariance matrix for the parameter likelihood conditioned on all but three parameters: process gain, process noise, and observation noise, each in the forward direction. Along two dimensions in this restricted parameter space, the covariance eigenvalues are quite large, indicating sloppy directions (***Gutenkunst et al., 2007***). This means that there is a plateau in the likelihood over parameters, with many distinct combinations of parameters giving quite similar likelihood, implying that we cannot infer these patterns with high confidence. However, for the remaining dimension, the covariance eigenvalue is much smaller, indicating that the data does reliably constrain this combination of parameters. This direction is interpretable as the weight given to the sensory evidence.

We then asked whether the different parameters for ASD and NT participants’ internal models could explain their differences in behavioral trajectories, which ***Figure 1*G** already showed contain enough information to differentiate between the ASD and NT groups. We attempted to categorize groups again using an SVM, but this time based on their inferred parameters and their uncertainties (see Methods Linear classification of participants’ parameters) rather than based on their complete trajectories.

The inferred parameters did successfully differentiate between ASD and NT groups (Kolmogorov-Smirnov test, *p* < 0.001, ***Figure 3*D**). The projection in ***Figure 3*D** shows the relative contribution of each parameter in distinguishing between groups. The most heavily weighted single parameter was the angular process gain (***Figure 3*F**). The assumed angular process noise variance and the subjective derivative action cost also contributed to discriminability between groups (***Figure 3*F**). Interestingly, when we shuffled the parameters between individuals within each group (i.e., removing parameter covariances), the SVM was unable to differentiate between the ASD and NT groups (***Figure 3*D** vs **E**; Kolmogorov-Smirnov test, *p* = 0.99), implying that the relationships between parameters were important. Thus, the uncertainties of inferred parameters should not be interpreted individually but must be considered in combination. In particular, due to data limitations, not all combinations of these parameters are trustworthy (See Methods Parameter degeneracy and reliability analysis). Thus we first identify the most reliable combinations, and then interpret them.

For the example participant shown in ***Figure 3*G**, the most reliable parameter combination is what we may call the ‘scaling of evidence,’ which is how much weight the participant’s updated belief (about their current position and velocity) gives to the joystick input *a*. On its own, different scalings result in updating posterior mean positions too much or too little relative to the ground truth, leading to under- or overshooting in the forward control dimension. However, combinations of parameters can obscure this effect, as one model property that contributes to overshooting can compensate for another model property that tends to undershoot. Alternatively, these model properties can manifest differently as functions of time or distance: for example, as shown in ***Figure 1*D**, participants tended to overshoot for near targets but undershoot for far targets ***Lakshminarasimhan et al. (2018***). In practice, the scaling of evidence has strong observable consequences for behavioral trajectories, which is why we can reliably estimate it.

Mathematically, the belief update actually depends on *two* terms: the intended joystick action *a*, and the sensory consequences *o* observed through optic flow and internal sensory noise. However, these experiments were abbreviated for our ASD patients, unlike our previous experiments (***Alefantis et al., 2022***; ***Kaushik J. Lakshminarasimhan, 2023***), so we did not include trials with action perturbations. Since the action and its sensory consequences were yoked together, any differences between *a* and *o* are therefore due to internal sensory noise, which we cannot directly measure. As a result, the two coefficients in the belief update Δ*b* = *c*_*a*_*a* + *c*_*o*_*o*, cannot be distinguished by our data; only their sum can be estimated. Note that both of these belief update weights *c*_*a*_ and *c*_*o*_ comprise multiple parameters, with their own influence on the weights derived from the parameter likelihood (see Methods Parameter degeneracy and reliability analysis). The reliable combination, leading to the overall scaling of evidence, is:

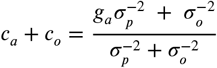

where *g*_*a*_, *σ*_*p*_, *σ*_*o*_, are process gain, process noise std, and observation noise std, respectively. For the participant shown in ***Figure 3*G**, the maximum likelihood parameters are gain *g*_*a*_= 0.7, process noise std *σ*_*p*_ = 0.6, and observation noise std *σ*_*o*_ = 1.0. The data is incompatible with modest *deviations* from these parameters, specifically for deviations along the combination process gain = 0.9, process noise std = 0.5, observation noise std = −0.1, since concerted changes along this parameter combination would change the behavioral bias from its measured value.

In contrast to this reliable parameter combination, there are some deviations that the data do not constrain. To illustrate this, we computed the covariance matrix for the three parameters above (process gain, process noise, and observation noise), now conditioned on all other parameters; we found that, in addition to the reliable dimension described above, the two remaining dimensions had much higher uncertainty (the “sloppy” directions (***Gutenkunst et al., 2007***), middle and right column in ***Figure 3*G**; Methods Parameter degeneracy and reliability analysis).

In the same way, we analyzed the uncertainty for the entire parameter set and identified several reliable dimensions (***Appendix 1 Figure 8***). Interpreting combinations of the entire parameter set is more challenging than for three parameters. In the following sections, we will focus our analysis on the particular parameter combinations that reliably differentiate ASD from NT.

### Individuals with ASD underestimate self-driven rotational velocity

We computed the reliable parameter combinations that most effectively differentiate ASD participants from the NT group. Since the 10-dimensional complete parameter combination is hard to interpret directly, and the correlation between sensory and motor parameters is low, we partitioned the parameters into two distinct sub-groups: those directly related to the sensory accumulation, and those directly related to motor preference, and interpreted them separately.

In the sensory parameter sub-group, we found that individuals with ASD have a lower ‘scaling of evidence’ and may ascribe a relatively lower weight to their actions in changing the world state. This could be a result of two different factors: (1) the process gain they assume, and (2) the relative reliability of their sensory observations versus their sensory predictions based on their current actions. Unfortunately, within the current abbreviated experimental design, these hypotheses cannot be distinguished. Accordingly, the joint log-likelihood surface shows a prominent ridge along many combinations of parameters that account equally for the data (***Figure 4*C**). While parameter combinations along the ridge cannot differentiate between the two groups, combinations orthogonal to the ridge do reliably distinguish ASD from NT.

**Figure 4.**
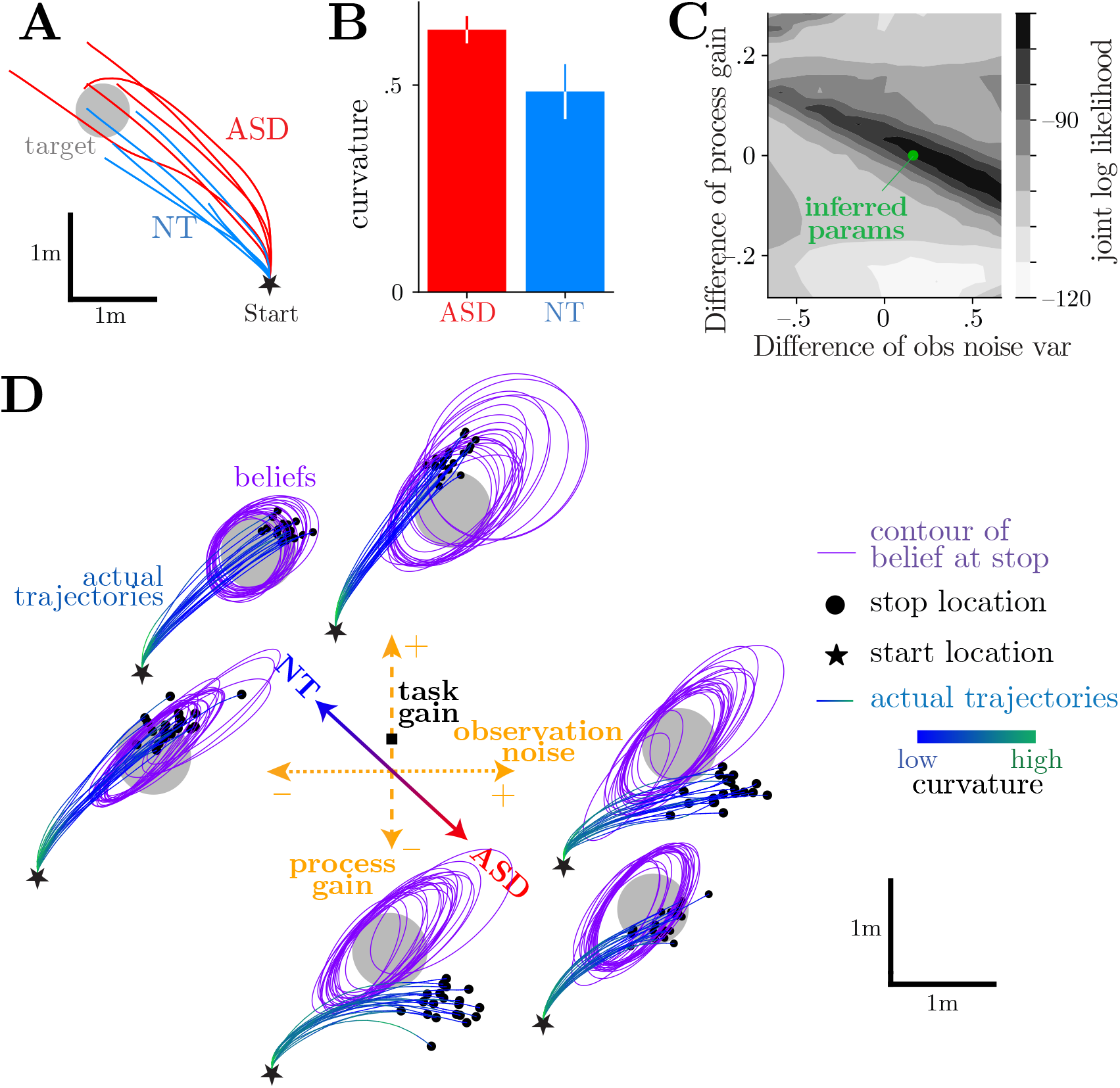
IRC reveals that ASD participants weigh their observations less. **A**. Example ASD and NT participant behaviors in one overhead plot. Black: target. **B**. ASD paths have a higher curvature (one tail *t*-test *p*-value = 0.04) for all side target trials. The error bar indicates the standard error of the mean. See precise definition of curvature in Methods Curvature calculation. **C**: Joint log-likelihood when varying 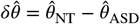. More than one combination of delta angular observation noise and delta angular gain can explain the different behavior of the two groups equally well. **D**: The model produces paths with high curvature like the ASD participants, whenever it underestimates its angular speed. This can happen due to a strong slow-speed prior that is given low weight because sensory observations are reliable, or due to a weak slow-speed prior that is nonetheless given high weight because sensory observations are unreliable. We are unable to identify the specific weight using currently available data, although experiments with control perturbations would have been able to distinguish between such options (see ***Appendix 1*** Additional model fitting validation). Purple ellipses: beliefs about the stopping position for each trial. Ellipses are centered on the best estimates, and the size and shape reflects uncertainties (represented by 2 standard deviations). Grey: target. Green-blue: actual trajectories, colored by curvature.

***Figure 4*D** visualizes the effects of changing these parameters. We choose the parameter vector halfway between the best parameters for ASD (***Figure 4*D** red axis) and NT participants (***Figure 4*D** blue axis). Then, we plot several trajectories for parameters that deviate from this mean, along various directions: toward smaller or larger process gain (***Figure 4*D**, vertical orange axis), or toward smaller or larger observation noise (horizontal orange axis). As shown by the inset illustrations, trajectories become more curved when either decreasing the magnitude of the process gain, or increasing the observation noise; both of these factors increase the relative weighting of the process gain relative to visual observations.

In the actual data, we observed similar behavioral difference pattern in the angular dimension when we examined trajectories for eccentric fireflies (i.e., targets that required more turning than the median, ***Figure 4*A**). On average, ASD participants’ trajectories had more exaggerated curvatures than did NT participants (*t*-test, *p* = 0.10; ***Figure 4*B**). This suggests that individuals with ASD acted as if they had to deflect the joystick more to achieve a given rotation. This results in higher curvature trajectories. (Having a higher cumulative angle turned does not contradict having the same end response angle from origin. See ***Appendix 1 Figure 4***.) This result aligns with our model prediction.

Together, these results demonstrate that IRC can reliably differentiate between the internal models of individuals with ASD from those of NT participants. The identified parameters suggested that participants with ASD perceived their self-driven rotational velocities differently from the NT participants, having a lower ‘scaling of evidence’ due to either lower process gain or a relatively stronger weight on the process gain than on visual observations. This explanation is supported by experimental data that individuals with ASD had exaggerated curvatures for eccentric targets.

### Individuals with ASD have a reduced subjective cost for actions

Among the complete parameter set, the only parameter that directly affects participants’ motor control is the subjective costs of acting. Specifically, we define a *derivative* action cost as the energy expenditure required to move the joystick from its current position. This is operationalized as the Total Squared Variation (TSV) of the trial joystick deflection.

In the IRC framework, this cost is scaled by a subjective parameter for each participant. Our analysis revealed that ASD participants showed a reduced subjective cost of acting compared to NT participants, implying differences in how they perceive their motor effort.

To illustrate how a reduced action cost can manifest in behavior, we simulated artificial agents that differ only in their angular action cost (***Figure 5*A–C**; low vs. high cost). These agents were presented with a visual perturbation from 1–2 sec (red region in ***Figure 5*A–C**). Their observations indicate they should adjust their motor output during or after the perturbation period to reach the target. As a control, we also instantiate an agent that does not receive a perturbation for which it must compensate (***Figure 5*C**). These simulations demonstrated that when actions are deemed low cost, agents may abruptly change their control dynamics during the perturbation (***Figure 5*A**, most prominently in forward velocity). Instead, when actions are perceived as high cost, agents compensate for the perturbation by distributing their control changes across many timesteps (***Figure 5*B**). It is ‘cheaper’ to maintain a small joystick deflection throughout the trial than to make a single abrupt, transient deflection.

**Figure 5.**
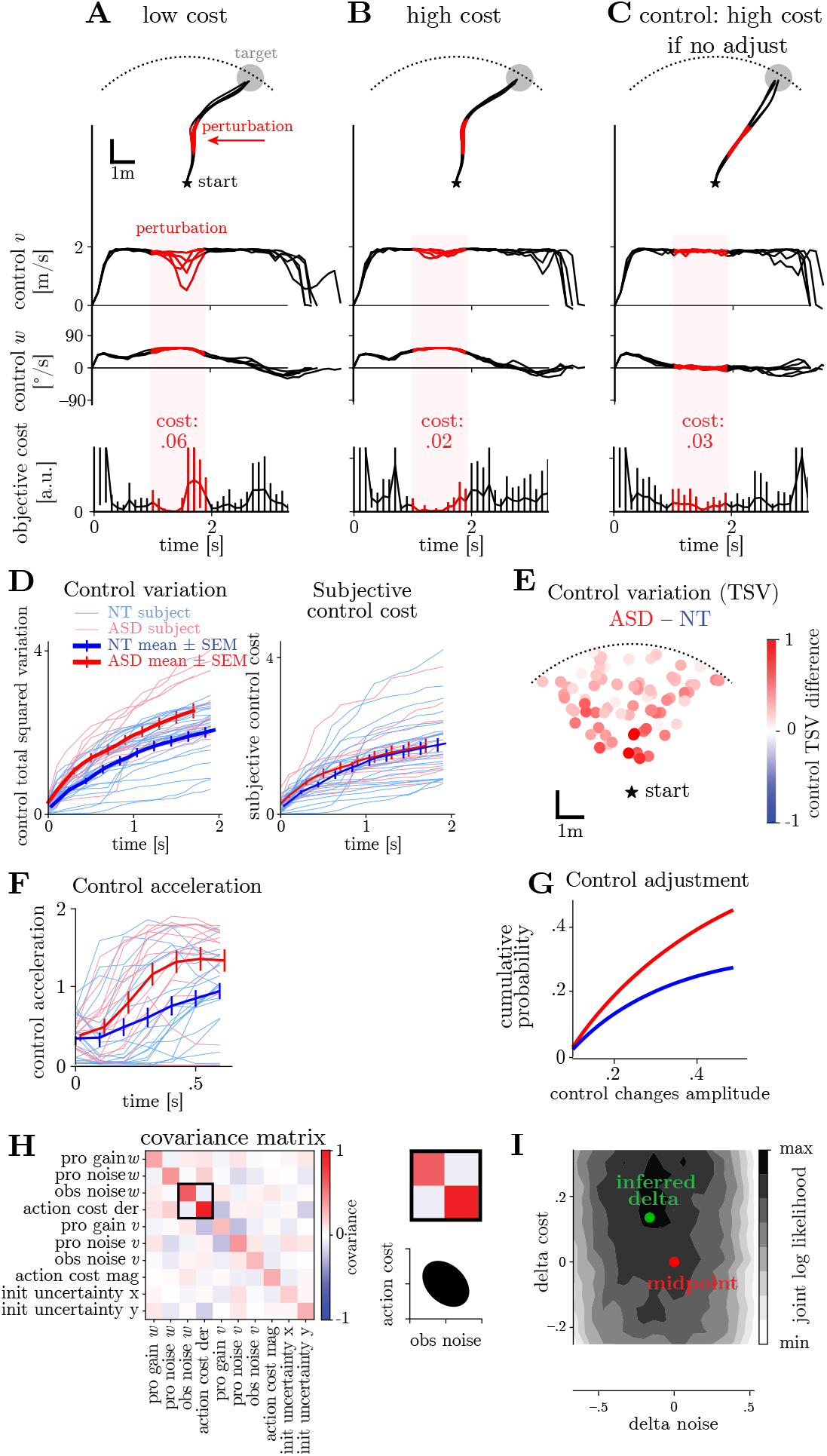
IRC can distinguish the effects of perceptual inference and motor costs. **A**. When the derivative action cost is low, the agent responds to the perceived process noise or perturbations by making an abrupt change in action. Black line: belief paths. Grey dot: the target. Red region: times when the agent receives a visual perturbation. The agent thinks he is ‘pushed’ leftward. **B**. When the derivative action cost is high, the optimal strategy is to distribute the adjustments into all time steps to avoid sudden changes in control, which is more costly. Compared to panel E (low-cost case), both forward and angular control curves (especially the forward control curve) are flatter when the angular cost is high. **C**. Control. When the control cost is high and the observation is not biased (no perturbation), the model does not need to make any adjustments. Both forward and angular control curves stay flat during the perturbation period. **D**. The accumulated control change and subjective control cost of ASD and NT participants navigating to the same example target. ASD participants have larger changes in their control. The inferred subjective action cost is similar across the two groups. **E**. The difference in ASD and NT group mean cumulative control change by all possible targets. ASD participants have higher control variation for all targets. **F**. At trial onset, ASD participants use greater angular acceleration, or, equivalently for velocity control, more change in their controls *da*_*ω*_/*dt*. **G**. Cumulative probability of acceleration amplitude. ASD participants change their controls more over the course of the trial. **H**. An example likelihood covariance matrix of the inferred parameters for one ASD subject. There is a weak anti-correlation between action cost and observation noise parameter. **I**. The joint log-likelihood of the two groups’ data when varying action cost and observation noise. There is only one global maximum thus one combination of action cost and observation noise parameter to explain the ASD behavior.

Consistent with these simulations, we found evidence from the actual data supporting that ASD have a lower subjective cost of acting. In other words, they use more costly actions to achieve the same control goal, compared to NT participants. In ***Figure 5*D** we plot both the TSV of the joystick control used by ASD and NT participants when navigating toward a given target, as well as their internally estimated (i.e., subjective) cost. The results demonstrate that, on average, individuals with ASD exert more control variance (TSV) than their NT counterparts (*t*-test *p* < 0.001, ***Figure 5*D**, left), yet do not perceive their actions as more costly than NT controls would (subjective cost; *t*-test *p* = 0.27, ***Figure 5*D**, right). This was true for all target locations (***Figure 5*E**), suggesting that the reduced subjective cost associated with actions present in ASD is a general phenomenon, and not associated with idiosyncrasies in navigating toward particularly easy or difficult (e.g., eccentric) targets.

Based on the lower subjective cost of actions in ASD (***Figure 5*D**), we predicted two behavioral differences. First, individuals with ASD should exhibit greater acceleration upon trial onset than NT participants. Second, ASD individuals should have larger control variations throughout the trial compared to NT participants. The experimental data supports both predictions, as presented in ***Figure 5*F**,**G**. Specifically, individuals with ASD exhibit greater acceleration upon target presentation (*t*-test *p* = 0.04) and a higher probability of making larger changes in joystick position compared to NT participants.

Given that the firefly task involves complex sensory and motor control interactions, we needed to determine whether differences in control preferences might be confounded with perceptual differences. Thus we examined whether the different subjective action costs for ASD and NT participants are dissociable from perception. For instance, one may avoid changing the current action to keep action costs low, or because of uncertainty associated with the world state and observations. To test this possibility, we examined the covariance matrix of inferred parameters. ***Figure 5*H** (example participant) shows both the full covariance matrix, and an inset focusing on the weak correlation between action costs and observation noise along the angular dimension. The mean correlation over all participants was only −0.03 ± 0.02 SEM. In turn, the joint log-likelihood of these two parameters in accounting for observed data demonstrates a single local maximum (***Figure 5*I**), not an extended ridge. This demonstrates that the estimated action costs were dissociable from observation noise, and thus reduced subjective action costs in ASD are not due to difference in perception, but action. As a result, reduced subjective action costs in ASD only caused ‘jerkiness’ in the trajectories (as seen in ***Figure 5*D** actual data, and ***Figure 5*AB** simulation data), not the exaggerated curvature observed in the earlier section.

## Discussion

Our results demonstrate that the framework of IRC was able to identify both an anomaly in closedloop perception in ASD (underestimated self-driven angular velocity), and dissimilarities in the subjective cost associated with movement (according to our model, ASD participants perceive their actions as less costly than their neurotypical counterparts do). These findings provide important insights into the underlying computational differences in sensorimotor control in individuals with ASD.

Individuals with ASD differ from their neurotypical (NT) counterparts in a host of domains (***American Psychiatric Association, 2013***; ***Robertson and Baron-Cohen, 2017***; ***Fournier et al., 2010***). Computational methods (e.g., ***Lawson et al. (2017***); ***Lieder et al. (2019***); ***Arthur et al. (2021***)) promise to establish links between seemingly disparate behaviors present in the condition. However, the field of computational psychiatry has so far largely focused on the study of perception (e.g., ***Noel et al. (2021***)) and disregarded the fact that perception and action form an inextricably closed loop. Here, we study the natural closed action-perception loop that exists in navigation. To do this, we apply the computational framework our group developed (IRC; ***Wu et al. (2020***); ***Kwon et al. (2020***)). In this approach, we first use reinforcement learning to build agents that can solve a broad family of related tasks, with parameters describing observations, states, actions, and subjective rewards. Then, from that family, we find the task whose solutions generate behaviors that most closely resembles the measured behaviors.

As a framework, IRC has important advantages over other comparable approaches, allowing for a flexible model for all aspects of the firefly task. Many related inverse models make more restrictive assumptions that make it harder to fit or interpret behavior. For example, some assume we know the observer’s dynamics ***Ziebart et al. (2008***); ***Choi and Kim (2011***); ***Babes et al. (2011***) or their objectives ***Dvijotham and Todorov (2010***); ***Schmitt et al. (2017***); others assume the observer fully observes their environmental state ***Ng et al. (2000***); ***Abbeel and Ng (2004***); ***Ziebart et al. (2008***); ***Dvijotham and Todorov (2010***); ***Finn et al. (2016***); ***Herman et al. (2016***); ***Reddy et al. (2018***). In contrast, IRC allows for unknown dynamics, objectives, and partial observability. A few studies in cognitive science use a similar approach to our method, allowing inversion of a POMDP but differing in their specific assumptions about the costs and controls, such as assuming a static task ***Houlsby et al. (2013***) or allowing only linear dynamics ***Straub and Rothkopf (2022***). IRC ***Kwon et al. (2020***); ***Wu et al. (2020***) better meets the needs of our task by allowing for a wider range of dynamic behaviors and mental models while retaining human-interpretabity. While we have certainly made some specific assumptions, like exponential discounting and stationary policies, that could be relaxed ***Schultheis et al. (2022***); ***Straub et al. (2023***), the current modeling formulation provides a strong match to our task.

One of the key advantages of IRC is that we can disentangle the sensory and motor properties by which ASD differs from NT, highlighted in our naturalistic firefly task. On the motor side, we found that individuals with ASD demonstrate a reduced subjective cost in executing actions. This manifests as more abrupt and more frequent changes in actions. Prior work has hinted toward this effect (***Mosconi et al., 2015***; ***Wang et al., 2015***) in a gripping task. When needing to maintain a steady grip force to match the required force prompted on screen, ASD participants showed more sudden and variable grip forces over time compared to NT. Our model ascribes both of these effects to a particular parameter, the lower subjective cost associated with changing actions.

On the sensory perception side, our results show that ASD individuals have a biased prediction gain and cannot predict the consequence of motor action as accurately as NT. This is consistent with behavior in grip tasks: when asked to generate a grip force to match a visual target shown on screen without immediate visual feedback, individuals with ASD consistently overshot the required force ***Wang et al. (2015***), suggesting deficits in accurately predicting the consequence of motor actions. ASD participants also maintain force without sensory feedback better than NT ***Shafer et al. (2021***), suggesting an underweighting of sensory evidence. Similarly, we observed ASD trajectories were more curved because the participants underestimated angular velocity predicted from their current actions. In accordance with our results, these findings suggest that ASD participants overweight internal predictions and underweight the external observation.

In our study, we found that ASD and NT participants had different perception of angular movement, but not forward movement. It is possible that our task setup cannot reveal the difference in the forward dimension due to a ceiling effect: the task may be relatively easy in the forward dimension, so both groups have similar, near-optimal performance. Only when the task is challenging enough, such as in the angular dimension, can we observe ASD’s distinct behavior and attribute it to their internal model deficit. Studies (***Manuel Vidal, 2009***; ***Paul A Warren, 2009***; ***M, 2000***) suggest that angular optic flow is indeed interpreted with greater uncertainty. This may be because in screen-based tasks like ours, the cue conflict between optic flow and vestibular input is stronger for rotation: The vestibular input is consistently active during rotation but adapts quickly to constant forward velocity. Moreover, in our experiment optic flow has less reliable information in the angular direction than in the forward direction (***Appendix 1 Figure 7***). Together, the greater cue conflict and weaker sensory signal should lead to greater uncertainty about angular movement, and thus more reliance on an internal model. The differences in ASD and NT internal models are then magnified in angular movements, differences that IRC can expose.

As mentioned in the results, there are two causes that could contribute to why ASD participants underestimate angular velocity: (1) they assume a slow speed prior ***Stocker and Simoncelli (2006***); ***Lakshminarasimhan et al. (2018***) that’s even slower than NT assume, and/or (2) they rely on their internal model velocity prediction from their current action more than NT participants do. Based only on our experiment, we are not able to identify how much each cause played a role in generating their velocity perception. However, building on findings from other studies, we can refine our interpretation. Our task includes an additional control component, making it a fully closed-loop system. Yet the belief update still uses classical Bayesian integration at each time:

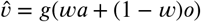

where *g* is gain (scaling introduced by the prior), *a* is action, and *o* is visual observation. Participants have a prior about velocity, which is combined with proprioceptive observations (translated into predicted velocity via the joystick action times the process gain) and visual observations (from the task optic flow) through optimal Bayesian integration to develop a velocity estimate. Evidence from prior research suggests that individuals with ASD perceive velocity equally precisely as NT participants when presented with fully coherent kinetograms (with limited dot lifetime, analogous to the optic flow in our task)***Catherine Manning (2013***), so we propose that ASD participants have similar levels of observation noise as NT. Evidence also suggests ASD participants have comparable accuracy in proprioception ***Christina T. Fuentes (2010***). In the context of the equation, this means ASD and NT have the same weights *w* and thus the same integrated sensory data (*wa* + (1 − *w*)*o*) in absence of a prior. Consequently, we propose that ASD participants possess a stronger prior for slower angular speeds, which results in their underestimated perception of angular velocity.

However, this interpretation assumes that sensory noises are identical between groups, a strong assumption, and attributes ASD distinct behavior solely to differences in their weightings. It is possible instead that ASD participants could have different sensory noises ***Simmons et al. (2009***); ***Zaidel et al. (2015***); ***Davis and Plaisted-Grant (2015***) (both visual and proprioceptive) than NT individuals. Additionally, sensory deficits in ASD participants could lead to biased observations, and/or biased proprioception. Accounting for these possibilities could yield different interpretations.

### Limitations and generalizations

Our study has important limitations. We were unable to determine the relative importance of the two mentioned above potential causes of the underestimation angular velocities in ASD. To address this, we need more experimental manipulations: specifically, it would be helpful to introduce a transient mismatch between actions and their effects on the environment ***Kaushik J. Lakshmi-narasimhan (2023***). When exposed to such perturbations, the participants behavior would reveal if they downweight or even ignore their prediction and instead rely more on their visual observations, or if they instead discount the sensory evidence and trust their internal model. We verified in simulation that this successfully resolves the ambiguity between prediction and observation (***Figure 2*E**). If we had more experimental time with this ASD patient population, this would be a high-priority addition to our experimental protocols.

Our explanation of ASD cognition is limited by the model we used. While no model can perfectly replicate the complexity of the human brain or produce indistinguishable artificial human behavior, there are important behavioral aspects that could be better captured by more complex models in future work. For instance, our current model assumes noiseless and cost-free internal representations and computations, yet in reality, belief may be noisy and suboptimal, and use energy costly spikes to encode information. Future models could account for how the brain balances the trade-off between representation cost and accuracy ***Castillo et al. (2024***). Finally, this study focuses on single-trial behavior, without accounting for fluctuations in participant behavior within a session, which could be influenced by factors such as fatigue. By incorporating these richer behavioral features into future models, we may be able to address more nuanced questions regarding ASD cognition.

Here we restricted the model-fitting exercise to well-learned behavior. Participants had *∼*150 trials of training prior to recording our experimental data, and had access to performance feedback during this training. Thus, during learning, we expect that participants’ understanding of the task changed gradually. Without explicitly modeling the learning process we would be unable to fit these changing behaviors. In contrast, by fitting behavior after extensive training with feedback, we elicited fairly accurate behavior, leading to relatively small discrepancies between the true and believed trajectories.

Naturally, we could have achieved more precise internal model estimations with additional data. Studies suggest that ASD has a large within-group variation (***Wozniak et al., 2016***), and we would ideally like to identify distributions of characteristics within this group. Our study incorporated relatively few subjects (ASD: *n* = 14; NT: *n* = 25) and few trials per subject (*∼*150), so cannot confidently draw any conclusions at the individual level. Given more data, we could potentially identify more structure within the autism spectrum. Lastly, we did not have access to detailed clinical profiles, which could have allowed us to correlate ASD symptoms and severity with behavioral profiles provided by IRC.

## Conclusion

This paper represents the first application of continuous IRC to behavioral data. Our results demonstrate the utility of this approach in characterizing complex, closed-loop, and continuous-time behavior based on a partially observable world. More generally, we expect this method will help advance our understanding of atypical and neurotypical brain function and provide computational targets for studying the neural mechanisms of perception, cognition, and action.

## Methods

### The firefly task

Human participants were tasked with “catching fireflies” in 3D virtual reality. The environment had no fixed landmarks and instead was composed solely of transient ground plane textural elements (***Figure 1*A**; size: base × height = 8.5 × 18.5 cm; density = 2.5 elements per m^2^). Participants used a joystick to control their virtual velocity in the 2D ground plane, move within the environment (maximum linear velocity = 2 m/s; max angular velocity = 90 degree/s), and stop at the firefly location. As participants moved, the ground plane elements correspondingly moved on the image, providing an optic flow signal indicating self-motion velocity. To estimate position from this evidence, participants had to continuously integrate the velocity (***Figure 1*A, B**). The target “fireflies” were presented for 1 s within an annulus sector with distances between 1 and 6 meters, and angles between ± 42.5 degrees from straight ahead (***Figure 1*C**). Upon stopping, participants received feedback about the trial outcome by a tone and an arrow pointing to the center of the firefly, colored green for trials rewarded (within 65 cm of the target) and red for unrewarded ones.

### Participants and behavior

This study involved 14 individuals with severe Autism Spectrum Disorder (ASD) and 25 neurotypical (NT) participants. Both ASD and NT participants had normal vision and motor function. Prior to the experiment, participants completed approximately 150 familiarization trials to become accustomed to the task and virtual environment. During experimental trials, participants had access to two sources of velocity information: optic flow and an internal model that mapped joystick position to virtual velocity. Participants were instructed to navigate towards a transiently visible firefly target using a joystick. The experiment consisted of two blocks of 150 trials each. In the first block, no feedback was provided about success or failure, to let participants learn the general controls during navigation. In the second block, feedback was given about the true target location. This paper focuses on data from the feedback portion of the experiment when participants were already familiar with the task and exhibited stable behavior. Joystick control trajectories were collected and down-sampled to 10 Hz for model fitting. For further experimental details please refer to (***Noel et al., 2020***).

### Inverse Rational Control framework

#### Forward model: generating behavior

We model rational human behavior in our task as an optimal agent solving a Partially Observable Markov Decision Process (POMDP). The objective parameters *ϕ* govern the world dynamics, including the state transition probability *T* (*s*_*t*+1_ |*s*_*t*_, *a*_*t*_; *ϕ*) between world states *s* given action *a*. In contrast, the subjective parameter vector *θ* represents the participant’s hidden assumptions and subjective preferences (Methods ***Table 1***). In rational control, as opposed to optimal control, these parameters *θ* may differ from the true world parameters *ϕ*.

Subjective parameters also govern the agent’s internal processes, such as the observation probability *O*(*o*_*t*_ |*s*_*t*_; *θ*) and the internal world state transition probability *T* (*s*_*t*+1_ |*s*_*t*_, *a*_*t*_; *θ*). In the experimentally defined firefly task, the state transition probabilities *T* (*s*_*t*+1_ *s*|_*t*_, *a*_*t*_; *ϕ*) and reward functions *R*(*s*_*t*_, *a*_*t*_, *s*_*t*+1_; *ϕ*) were deterministic. We modeled the participant’s observations as linear functions of the state with additive Gaussian noise, *o*_*t*_ = *Hs*_*t*_ + *η*_obs_, where the matrix *H* = [0 1] allows direct observation only of the velocity but not the position, and *η*_obs_ represents observation noise with covariance 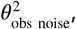, leading to a normal observation probability 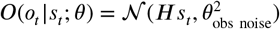.

Since the states are only partially observable by the agent, the agent cannot make decisions based directly on states but must instead use its beliefs and the anticipated transitions between them. By incorporating this uncertainty, the internally assumed state transition probability leads to *belief* transition probabilities 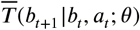 that account for their actions not their sensory observations. Parameters of this transition probability will incorporate subjective assumptions of the parameters in *θ* specifying the process gain and process noise. The transition and observation probabilities are then combined by Bayesian inference into a belief update probability *P* (*b*_*t*+1_ |*b*_*t*_, *a*_*t*_, *o*_*t*+1_; *θ*) accounting for both actions and observations. We modeled this process as an extended Kalman filter (***Kalman, 1960***), which weighs predictions and observation by their relative reliabilities. Finally, the beliefs determine a rational policy *π*(*a*_*t*_ |*b*_*t*_; *θ*) that is optimized under these transitions and the subjective expected reward 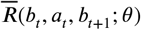, which is the state reward *R*(*s*_*t*_) averaged under the state posterior *p*(*s*_*t*_ |*b*_*t*_) represented by the belief *b*. Finally, over the continuous space of parameters *θ* we estimated the rational policy family Π(*a*_*t*_ |*b*_*t*_; *θ*) using the reinforcement learning algorithm TD3 (***Fujimoto et al., 2018***).

#### Model parameters

#### Inverse model: fitting the latent parameters

##### Likelihood calculation

To infer a participant’s latent subjective parameters, we invert the forward model and find which particular internal model and parameter vector best explains the participant’s behavior. Mathematically, we evaluate the goodness of fit for the estimated parameter *θ* using the complete data likelihood function and the Markov property of the POMDP,

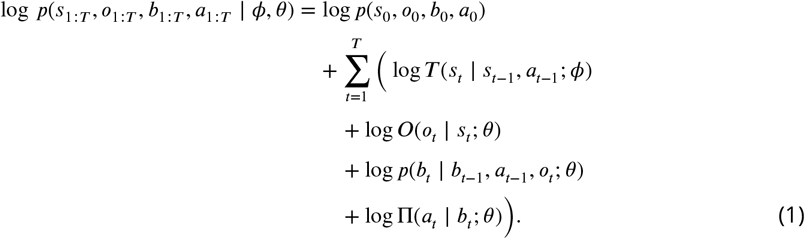

##### Maximizing the Log-Likelihood

This complete data likelihood depends on quantities we cannot observe, so we marginalize it over the latent observations *o*_1_ and beliefs *b*_*1*_ to obtain the observed data likelihood *p*(*s*_1:*T*_, *a*_1:*T*_ | *ϕ, θ*). This is done by sampling observations for each state *s*_*t*_ presented to the participants, drawing *L* samples based on the observation noise variance specified by *θ*. For our model, the belief update is deterministic given these observations, so there is no need to marginalize over the beliefs through additional sampling. The likelihood for one observation sample and its resultant belief is

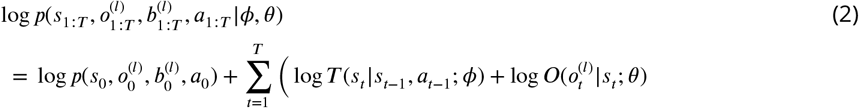

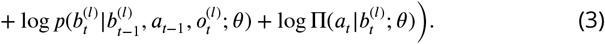

where *l* stands for a sample of the entire latent variable trajectory. Note that the only terms depending on the participant’s parameters *θ* are the latent observations’ probabilities, belief dynamics, and policy. Since we are optimizing over *θ*, all other terms independent of *θ* are irrelevant and can be neglected. After simplification, the best estimate of the participant’s parameters, 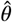, is:

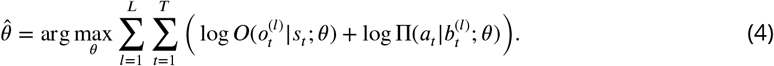

To solve this optimization problem, we used the Covariance Matrix Adaptation Evolution Strategy (CMA-ES (***Hansen, 2023***)). This method defines a population Θ of *N* potential solutions *θ*_*n*_ and allows that population to evolve toward improving the objective function. As with all optimization problems, there is a trade-off between exploration and exploitation. In this specific case, the likelihood surface in the parameter space is highly influenced by stochasticity induced by sampling, making gradient-based methods less reliable. CMA-ES is well-suited for this problem, and also provides uncertainty in the final estimate 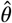.

The CMA-ES algorithm begins by initializing a starting population of *N* candidate parameter vectors *θ*, and estimating the log-likelihood of each *θ*_*n*_ in the population. We begin the optimization by sampling from the entire parameter space, with the mean centered on the mean of the entire parameter range and a wide isotropic covariance covering each parameter’s range within one standard deviation. CMA-ES then iteratively updates the mean and covariance of the population using rank-*µ* updates (***Hansen, 2023***). We stop iterating when all diagonal entries of Cov Θ are smaller than 0.005 of the original initialization. This corresponds to constraining the standard deviation of the forward gain parameter estimate to 0.1 m/s.

The IRC algorithm using CMA-ES can be summarized below:

###### Algorithm 1

IRC algorithm optimized using CMA-ES

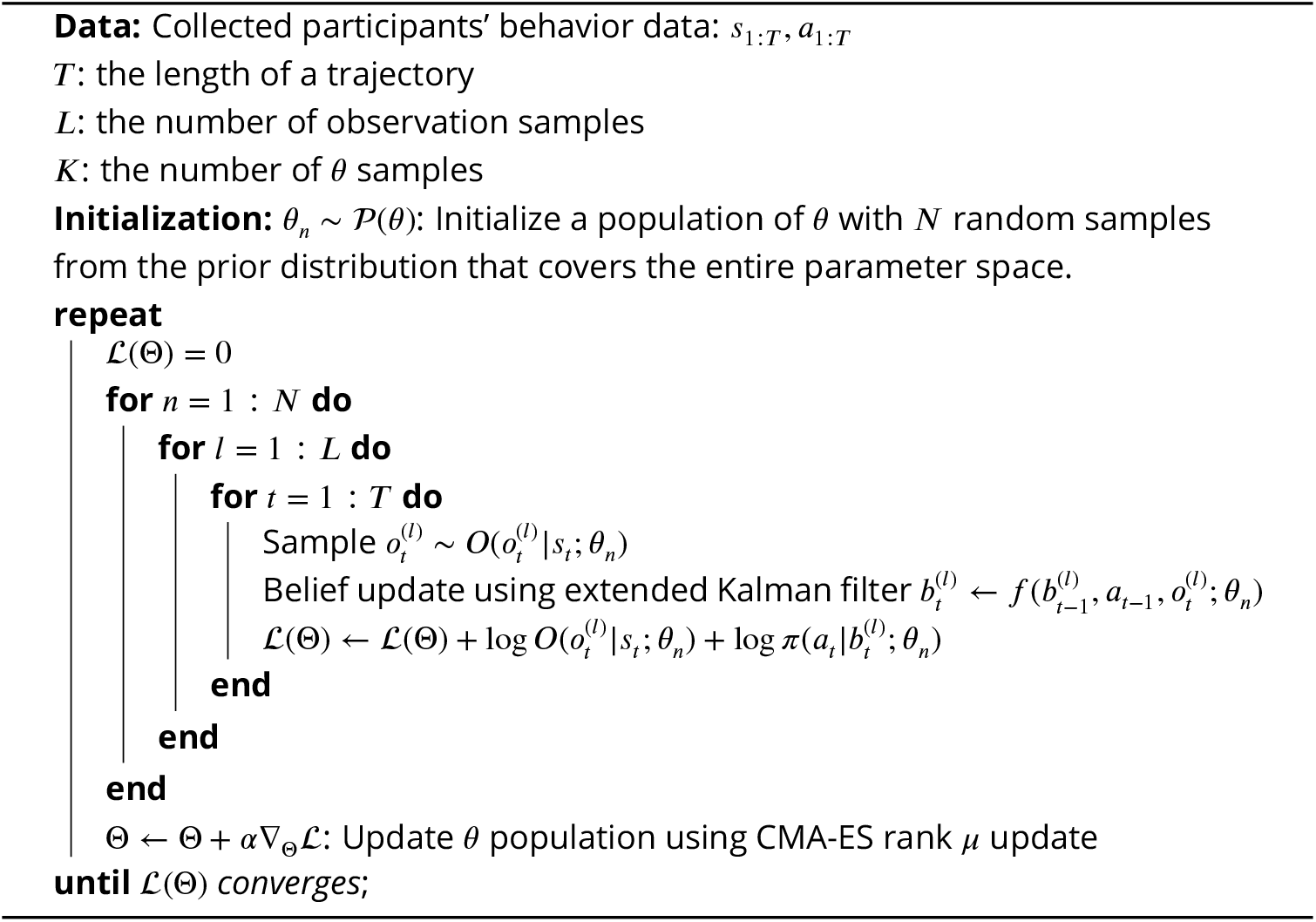

#### Validation of the IRC on simulation data

We tested this IRC approach over multiple runs using simulated data, leading us to conclude that the IRC was stable and this gives confidence that we can find global minima. Previously, we had demonstrated (***Kwon et al., 2020***) our ability to simulate and solve the inverse problem in a simpler context. In this study, we reproduced the result in a more complex simulation with additional constraints, such as multiple types of action costs and process perturbations (***Alefantis et al., 2022***; ***Kaushik J. Lakshminarasimhan, 2023***).

Specifically, we measure 1000 behavioral trajectories of an agent whose subjective parameters *θ*^*\**^ were randomly sampled from the allowed parameter range. Out of the 1000 trials, half used the same experimental task parameters as the participants used, and in the other half of the trials, we generated random perturbations similar to our previous studies (***Alefantis et al., 2022***; ***Kaushik J. Lakshminarasimhan, 2023***): we added both forward and angular velocities to the intended controls, within a smooth 1-second-wide bell-shaped temporal profile of standard deviation of 0.2 s, and uniformly-distributed random amplitudes between ±2 m/s for forward velocity perturbations, and between ±120°/sec for angular velocity. We then applied the IRC algorithm to infer the agent’s latent parameter *θ*. The result is shown in ***Figure 3*E**, where our best estimate of 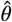 and its uncertainty is represented by an elliptical confidence region of one standard deviation (yellow), and the ground truth *θ*^*\**^ is shown as a white star.

### Linear classification of participants’ parameters

To distinguish categories of IRC parameters between ASD and NT groups, we trained a linear SVM classifier on their inferred parameters. For each participant we drew 50 parameter samples from a truncated Gaussian distribution with inferred parameter mean *µ*, covariance Σ, and support within the valid parameter range. Each sample was labeled as ASD or NT according to the participant’s group and used to train the SVM classifier. To visualize the classification results in ***Figure 3***D and E, we projected the samples onto the most discriminative axis identified by the SVM.

### Parameter degeneracy and reliability analysis

To quantify the reliability of inferred parameters, we computed the covariance matrix (inverse Hessian) of the parameter likelihood at its peak, as shown in ***Appendix 1 Figure 8***A. The (marginal) confidence interval for each inferred parameter is described by the diagonal values of the covariance matrix. However, the parameters have complex interactions, so it is helpful to examine the eigen-decomposition of this covariance matrix. Its eigenvalues reflect the confidence (variance) in estimating the parameter combinations along each corresponding eigenvector (***Appendix 1 Figure 8***B and ***Figure 3*H** for an easier-to-interpret conditional case): lower eigenvalues reflect narrower likelihood, and thus tighter constraints on that combination of parameters. Out of all inferred internal model parameters, the most reliable parameter combination is the eigenvector with the smallest eigenvalue (shown in ***Appendix 1 Figure 8***B, leftmost column).

### Comparing ASD vs NT trajectories

When comparing trajectories from ASD and NT participants, since we do not have exact repeats for the same target locations, we need to identify similar targets for comparison. For each target position on the map, we identify the three nearest target positions from both ASD and NT participants to compare. We limit this to three nearest targets because including more would include targets at a distance, potentially compromising the validity of our comparisons.

### Curvature calculation

The participant’s path is a set of points (*x*(*t*), *y*(*t*)). The curvature of this trajectory is

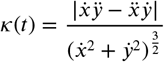

where the 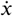,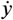 and 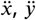 are the first and second temporal derivatives of *x*(*t*) and *y*(*t*), approximated by finite differences. We quantify a trajectory’s overall curvature as the temporal mean of *k*.

## Data Availability

Data is available at https://osf.io/chtjb/. Code for data analysis and figure generation is available at github.com/yizhoucc/firefly_plots.

## Acknowledgments

This work was supported by NIH grant CRCNS 1R01 NS120407-01 to XP and DEA, by grant SFI-AN-NC-SCN-00007276-10 to XP, JN, and DEA from Simons Foundation’s SCENE program, and by grant SFI-AN-AR-Pilot-00008952 to JN from the Simons Foundation’s SFARI program.

## Appendix 1

### Additional participant behavior

**Appendix 1 Figure 1.**
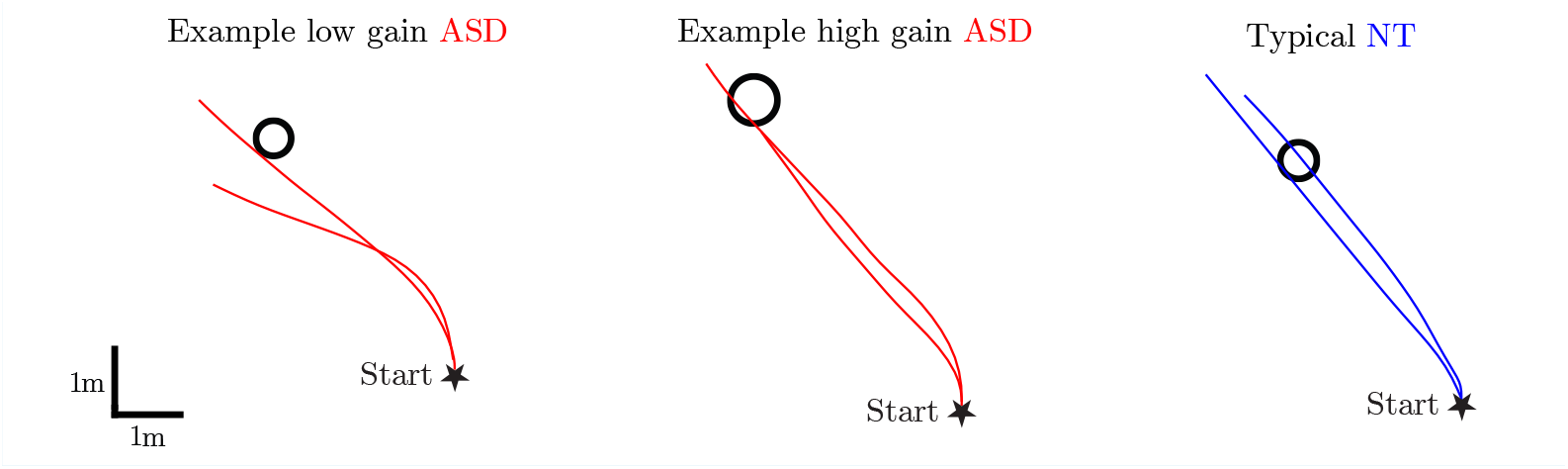
Example participant paths for the same target. ASD participants’ trajectories are more curved than NT participants’.

**Appendix 1 Figure 2.**
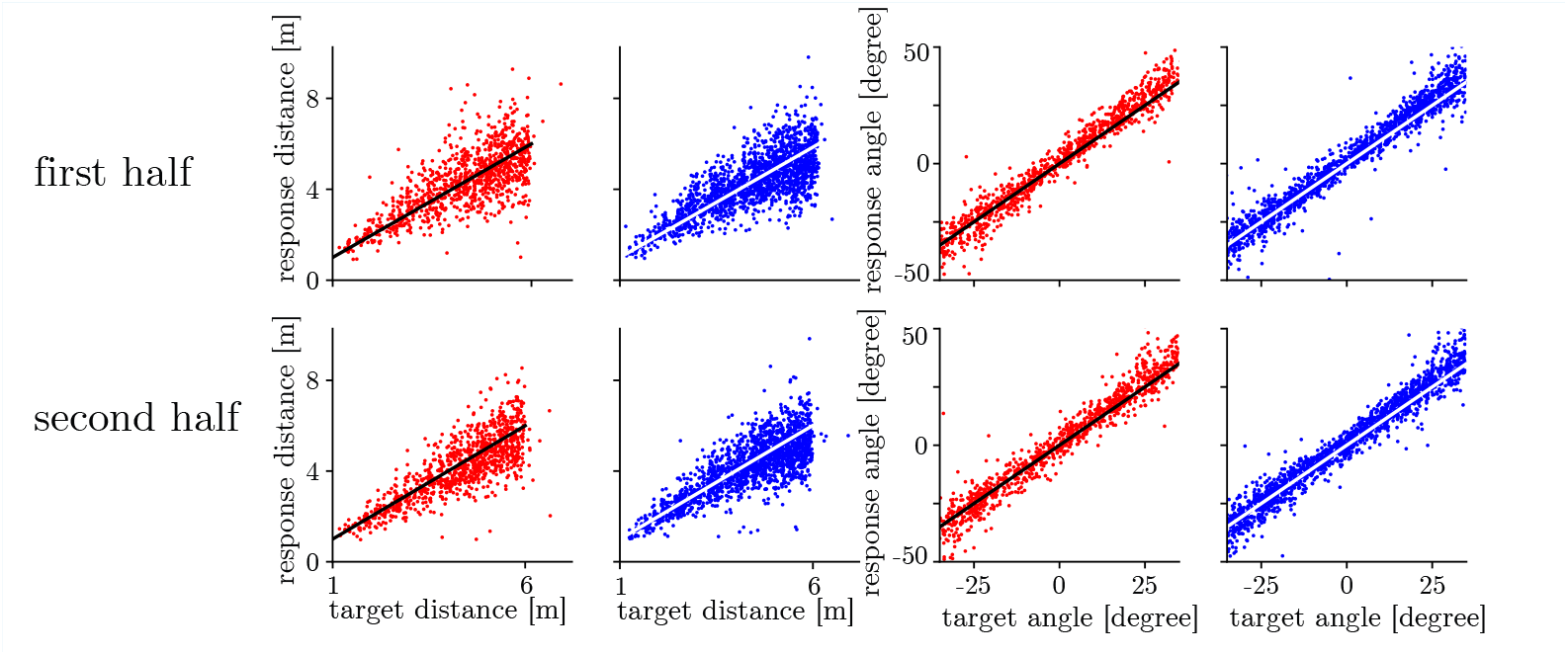
There was no significant improvement in performance between the first and last halves of the participant’s trials, indicating that no substantial learning occurred.

**Appendix 1 Figure 3.**
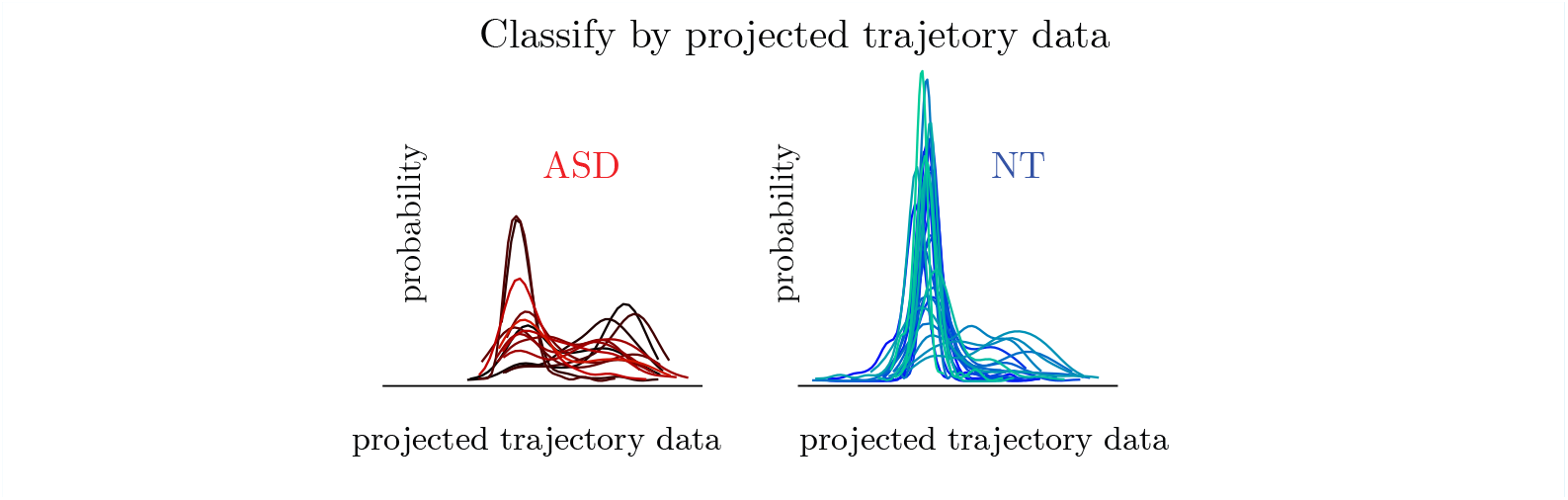
The distribution of ASD and NT trajectory data for each participant, projected onto the most separable axis identified by SVM. Left: ASD. Right: NT.

**Appendix 1 Figure 4.**
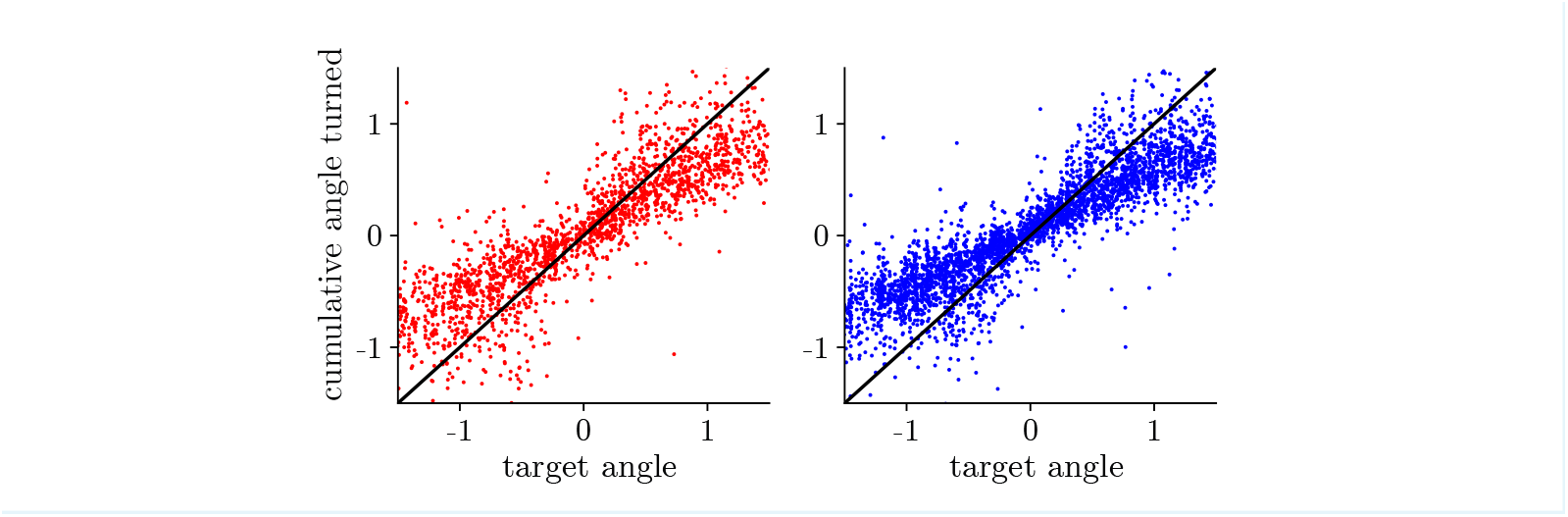
The ASD participants show a significantly higher cumulative angle turned than the NT participants. ASD slope: 0.64, SEM=0.01; NT slope: 0.57, SEM=0.01. *t*-test *p* value < 0.05. ASD participants prefer to reach the target with more curved trajectories. This is evidenced by their higher cumulative angle turned, while maintaining the same final response angle from the origin (***Figure 1*E**)

### Additional model fitting validation

**Appendix 1 Figure 5.**
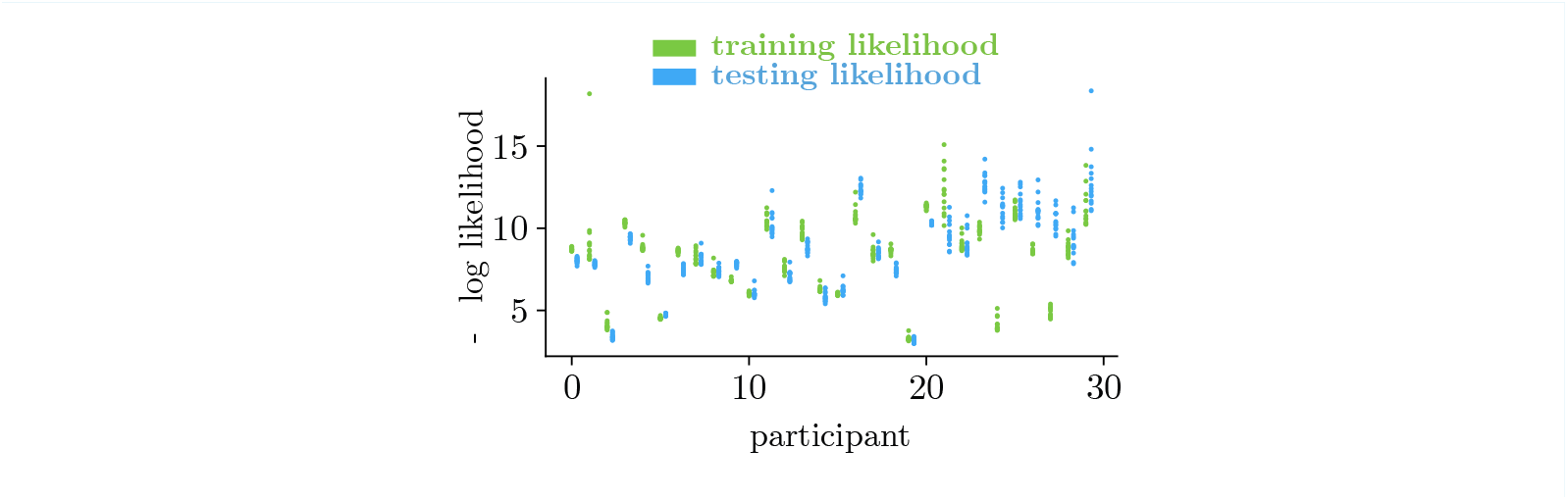
We cross-validated the inferred model for each participant on a held-out test dataset to assess out-of-distribution generalization. 75% of trials, those with shortest target distances, were allocated to the training set; the remaining 25% of trials, those with the longest target distances, were reserved for testing. We considered the model to fit well if it could accurately predict the participant’s behavior on these test trials. The log-likelihood, used as the evaluation metric, was comparable on both the training and testing sets. (mean difference = –1.19, not significantly different by *t*-test with *p* = 0.26)

**Appendix 1 Figure 6.**
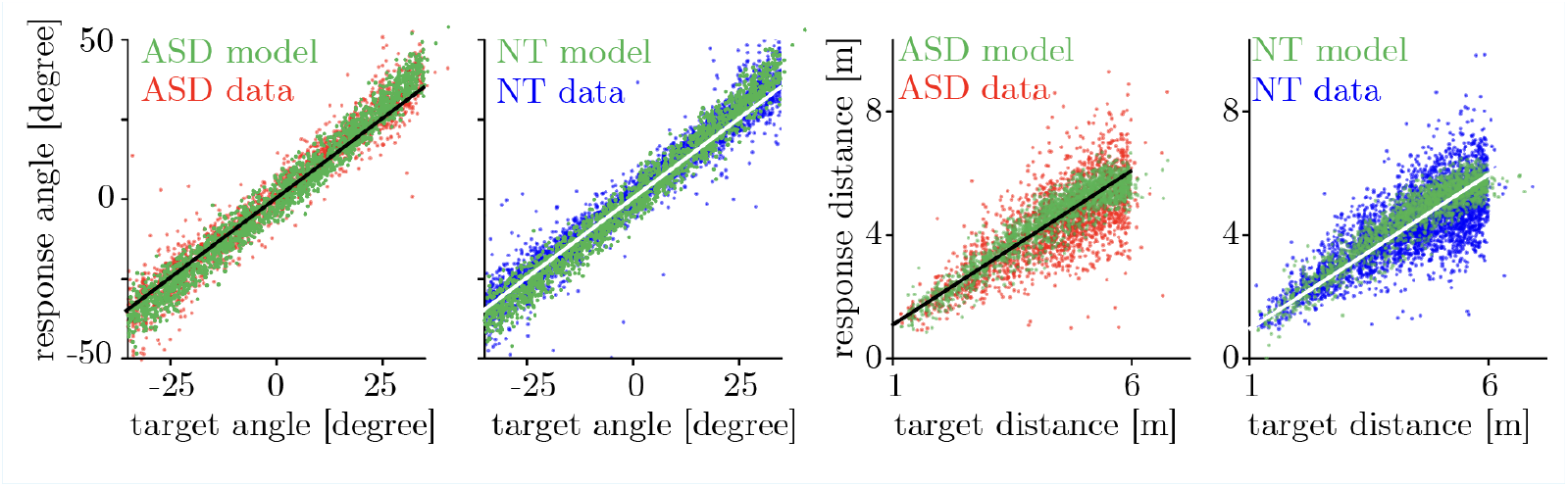
Comparison of model and actual data for end-point responses. The model accurately captures the bias and variance of angular responses, though it exhibits less variability in the final responses in the forward dimension than the actual data. For reference, black and white diagonal lines have a slope of 1.

### Simulation of the optic flow information

**Appendix 1 Figure 7.**
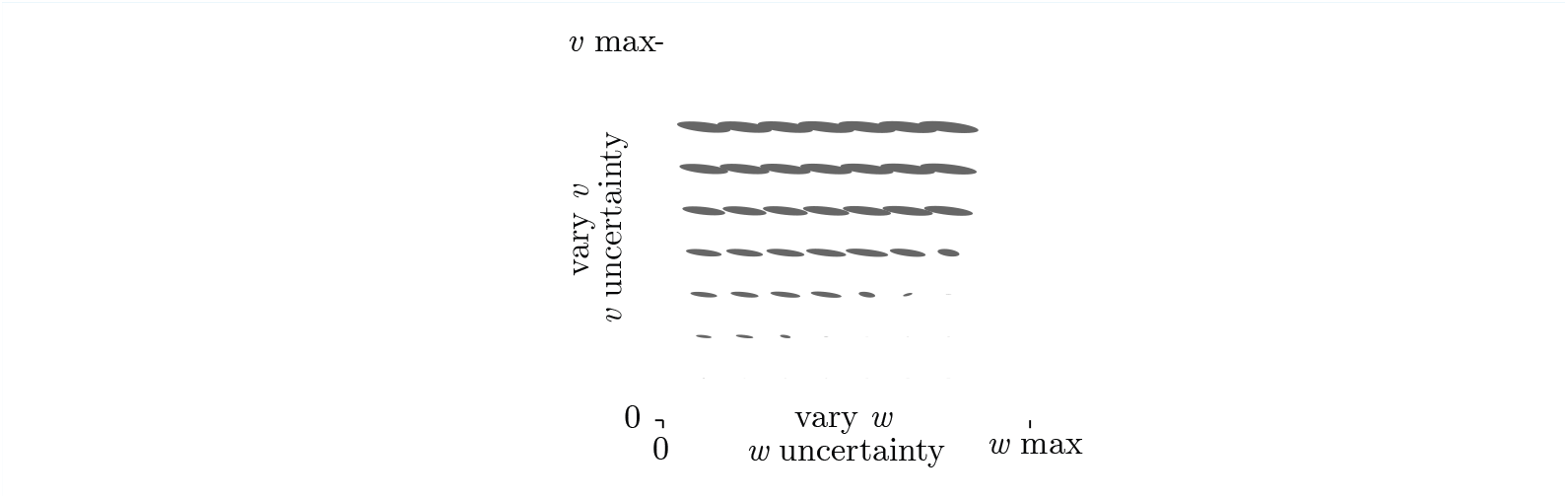
An ideal observer’s uncertainty about forward (*v*) and angular (*w*) self-motion, estimated from on-screen observations, with covariances depicted as grey ellipses representing one standard deviation per second and are scaled relative to the maximum forward velocity (2 m/s) and angular velocity (90°/s). First, we modeled the uncertainty in on-screen observations by assuming it follows Weber’s Law, such that the uncertainty, measured by standard deviation, increases proportionally as the optic flow triangle size decreases and on-screen velocity increases. We then used a geometric model of the task to transform the on-screen uncertainty to virtual world uncertainty. The geometric model assumes a virtual player height of 10 cm, a visual field of view of 90 degrees, and a maximum optic flow distance of 5 m, which corresponds to the actual task. Based on these parameters, we calculated the Jacobian matrix from the geometric model and used it to transform the on-screen covariance matrix into the 3D world. The resulting 3D world covariance matrix represents the uncertainty about the current motion, given the on-screen optic flow. The results demonstrate that within our virtual environment, optic flow provided more information about forward movement than angular movement.

### Forward model details

**Appendix 1 Table 1.** Hyperparameters in forward model reinforcement learning training.

| Hyperparameters | Value | Hyperparameters | Value |
| --- | --- | --- | --- |
| learning rate $r$ | $7 \times 10^{-4}$ | batch size | 512 |
| replay buffer size | $1 \times 10^6$ | discounting factor $\gamma$ | 0.96 |
| training frequency (per timestep) | 4 | action noise | $\mathcal{N}(0, \frac{1}{2} \mathbb{I})$ |
| training policy delay | 6 | random seed | 42 |
| policy network layer widths | [21,64,64,2] | layer activation function | ReLU |
| critic network layer widths | [23,64,64,1] | output activation function | tanh |
| target policy noise | 0.2 | target noise clip | 0.5 |
| tau | $5 \times 10^{-3}$ | | |

The agent is trained using a curriculum that progressively introduces tasks with increasing levels of uncertainty, from low to high. We also incorporate action costs to ensure that the agent’s actions are practical in real-world situations. The training sessions are structured to allow the agent to optimize for the entire session and overall reward rate, rather than focusing on each individual trial. As a result, the agent may choose to skip less valuable trials in favor of encountering more rewarding ones.

### Inverse model details

**Appendix 1 Table 2.** Hyperparameters in inverse model fitting.

| Hyperparameters | Value |
| --- | --- |
| CMA-ES population size | 14 |
| number of trial samples | 5 |

The learning rate in the inverse model fitting is automatically determined by the CMA-ES algorithm, which takes into account the population size, the current population, the best-fit subpopulation’s mean and covariance matrix, and the number of dimensions.

The CMA-ES method samples a population of *θ* from a Gaussian distribution Θ and optimizes this distribution. However, when Θ is near the boundary of the parameter space, resampling is required to ensure that the sampled *θ* values remain within the valid range. To prevent hard clipping of *θ*, which could result in a skewed distribution with excessive density near the boundary, we resample until we obtain valid *θ*_*k*_ values.

### Inferred parameter distributions for ASD and NT groups

**Appendix 1 Table 3.** Inferred parameter distributions for both groups, as plotted in ***Figure 3*C**. a.u. stands for arbitrary units.

| Parameter $\theta$ | | ASD mean±SEM | NT mean±SEM |
| --- | --- | --- | --- |
| Process gain | (forward) | 164±0.78 cm/s | 150±0.78 cm/s |
| | (angular) | 76±0.14 °/s | 81±0.17 °/s |
| Process noise | (forward) | 102±0.68 cm/s | 100±0.58 cm/s |
| | (angular) | 12±0.11 °/s | 12±0.11 °/s |
| Observation noise | (forward) | 179±0.89 cm/s | 161±0.64 cm/s |
| | (angular) | 44±0.16 °/s | 46±0.24 °/s |
| Initial uncertainty | (forward) | 76±0.71 cm | 49±0.53 cm |
| | (angular) | 34±0.39 cm | 33±0.38 cm |
| Action cost | (derivative) | 0.49±0.01 a.u. | 0.65±0.01 a.u. |
| | (magnitude) | 0.98±0.01 a.u. | 1.01±0.01 a.u. |

### Parameter degeneracy and reliability analysis

**Appendix 1 Figure 8.**
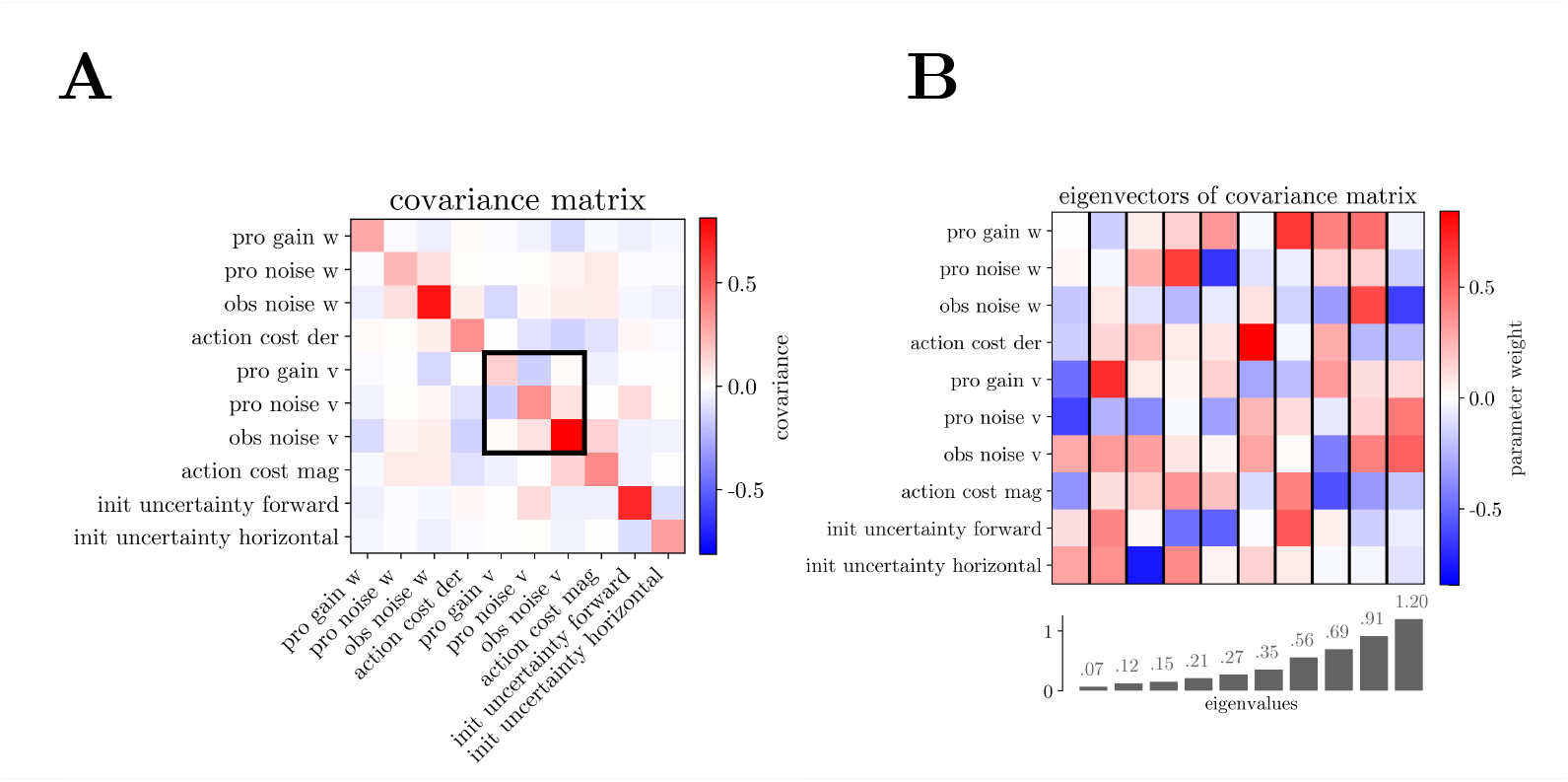
Covariance matrix for the parameter likelihood of an example ASD participant and its Eigendecomposition. **A**. Multivariate confidence regions for parameters are determined by the covariance matrix of the parameter likelihood (***Press, 2007***). Here we showed a covariance matrix for one example ASD participant. The 3×3 square outlined in black indicates the parameters for process gain, process noise, and observation noise, each in the forward direction. We took a conditional covariance matrix on all but these 3 parameters in ***Figure 3*G** for an easier-to-interpret close examination. **B**. Eigendecomposition (as described in Methods Parameter degeneracy and reliability analysis) of the covariance matrix in panel A. The eigenvalues reflect the confidence (variance) in estimating the parameter combinations along each corresponding eigenvector. Each column depicts one eigenvector, with colors corresponding to the contribution of each parameter. Below each column, the eigenvalues reflect the widths (variance) of the likelihood along the corresponding directions in parameter space

Out of all inferred internal model parameters, the most reliable parameter combination is the eigenvector of the covariance with the smallest eigenvalue (shown in panel **B**, leftmost column). This eigenvector places the greatest weight on the forward gain, forward process noise, and forward observation noise, as well as action cost and initial uncertainty.

On the other hand, other combinations of those same parameters are not identifiable. This non-identifiability occurs when distinct parameter values produce near-identical belief updates. According to Bayesian updating, the belief dynamics combines the prediction from the action *a* with an observation *o*, weighted by their precisions (inverse variances):

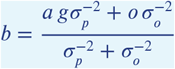

where the three parameters *g*, 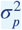, and 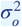 are the process gain, process noise variance, and observation noise variance, respectively. Here we cannot differentiate between the intended action and the observed optic flow, because in our experiment these only differ due to sensory noise, which we cannot directly measure here. (In the Discussion, we consider approaches that remedy this in future experiments by adding experimentally measurable observation noise.) Consequently, we cannot fully disambiguate the individual parameters. Nonetheless, we can identify the total combined weight on these optic flow components because that measurably affects the scale of the participant’s control and the resultant undershooting. This aligns with our result shown in ***Figure 3*G**.

